# IGP receptors act as redox sensors to mediate broad-spectrum disease resistance induced by cellooligomers

**DOI:** 10.1101/2025.10.05.680433

**Authors:** Guangzheng Sun, Pei Wei, Jingjing Dai, Jinyu He, Xuan Mi, Xinrui Li, Qinsheng Zhu, Ning Xu, Zhichao Zhang, Liangyu Guo, Yifei Yang, Mingzhu Fan, Yan Ren, Feng Chen, Ming Chang, Zhifu Han, Aolin Jia, Xiao-Quan Wang, Yuanchao Wang, Yu Xiao

**Author notes:** These authors contributed equally (Guangzheng Sun, Pei Wei, Jingjing Dai, Jinyu He).

## Abstract

Cys-oxidation is a well-established post-translational regulator of intracellular protein function, yet its role in extracellular receptor-mediated signaling remains poorly understood. Here, we investigate the LRR-Malectin receptor-like kinase IGP1, a key receptor for plant cell wall damage-associated molecular patterns (DAMPs). We demonstrate that IGP1 directly recognizes cellooligomers with a degree of polymerization ≥ 3 via a conserved binding groove in its LRR domain, not the malectin domain as anticipated. Strikingly, a pair of cysteines within the malectin domain functions as a redox sensor, undergoing hydrogen peroxide-induced oxidation that sterically reduces ligand binding affinity and attenuates immune signaling. Phylogenetic analyses reveal that IGP originated after the terrestrialization of plants, and is an evolutionarily conserved redox sensor in land plants. Functional studies in Arabidopsis (*Arabidopsis thaliana*), wheat (*Triticum aestivum*), and soybean (*Glycine max*) confirm that the IGP-cellooligomer module confers broad-spectrum disease resistance. Our work uncovers a unique mechanism where ligand perception and redox sensing are integrated into a single receptor to regulate DAMP-triggered immunity.

## Introduction

Signal perception through cell surface receptors constitutes a fundamental biological process across living organisms. In plants, this critical function is primarily mediated by receptor-(like) kinases (RLK/RKs) embedded in the extracellular matrix. Characterized by their highly diverse extracellular ligand-binding domains, RLKs are categorized into several distinct families, including leucine-rich repeat (LRR)-RLK, lysin motif (LysM)-RLK, *Catharanthus roseus* RLK1-like (CrRLK1L), and Lectin-RLK, etc. Recent studies have identified LRR-Malectin RLKs (previously classified as LRR-RLK VIII-2 subfamily) (*1–3*) as a unique receptor class featuring dual ligand-binding domains: LRR domains and a malectin domain. These receptors regulate multifarious physiological processes ranging from development (*4*), reproduction (*5, 6*), nutrient stress (*7*) and immunity (*8–10*).

A number of RLKs function as pattern recognition receptors (PRRs) detecting both damage-associated molecular patterns (DAMPs) from host plants and pathogen-associated molecular patterns (PAMPs) from invaders to induce pattern-triggered immunity (PTI) (*11*). The plant cell wall, a highly complexed carbohydrate matrix comprising cellulose, hemi-celluloses and pectins, acts as critical and the first line of defense against pathogen. To breach this barrier, pathogens secret various cell wall-degrading enzymes, triggering the release of various DAMPs: oligogalacturonides (OGs) (*12, 13*), cellooligomers (*14–16*) and oligosaccharides stemmed from mixed-linked glucans (MLGs) (*17*), xyloglucans (*18*), xylans (*19*) and mannans (*20*). While LysM-RLKs are well-established receptors for microbial chitooligosaccharides and its derivatives (*21*), the recognition mechanisms for plant-derived cell wall oligosaccharides as DAMPs remain poorly understood. Notably, the proposed Arabidopsis OG receptors (wall-associated kinases, WAKs) (*22*), were recently reported to be not required for OG-induced immunity (*23*), highlighting critical knowledge gaps in plant cell wall DAMP perception.

The malectin domain, originally characterized as an endoplasmic reticulum-localized glycan-binding protein MALECTIN that recognizes Glc₂-high-mannose N-glycans during the early steps of protein N-glycosylation in animals (*24*), has emerged as a key candidate for oligosaccharide perception in plants. Several malectin RLKs have been shown to demonstrate potential oligosaccharide-binding capabilities. CrRLK1L: FERONIA (FER) interact with rapid alkalinization factors (RALFs) (*25*) and pectin/OGs (*26, 27*). Malectin-LRR RLK: symbiosis receptor kinase (SYMRK) (*28, 29*) cooperates with nod factor receptors in symbiosis signaling. LRR-Malectin RLK: impaired in glycan perception 1 (IGP1, also known as cellooligomer-receptor kinase 1, CORK1) and IGP2/3,4 mediate perception of cellooligomers (*14, 15*), MLGs (*14*) and XYL4 (*30*) to trigger PTI response. However, despite these advances, the precise mechanism of malectin RLKs interaction with cell wall oligosaccharides remain less well elucidated.

Reactive oxygen species (ROS) and nitric oxide (NO) serve as crucial signaling molecules both intracellularly and extracellularly. The thiol group enables cysteine residues to function as redox switches, sensing environmental ROS/NO levels and undergoing reversible oxidation and reduction. These post-translational modifications alter protein structures, thereby modulating a broad range of biological processes. While such redox-mediated regulation has been extensively documented for intracellular processes recently (*31–37*), its role in the extracellular space remains poorly understood. To date, only HPCA1, an LRR-RLK, has been identified as a hydrogen peroxide sensor (*38*).

In this study, we elucidate the mechanism of cellooligomer perception by an LRR-Malectin RLK IGP1. Integrated phylogenetic, biochemical, structural, genetic, and pathological analyses reveal that this evolutionarily conserved IGP receptor possesses a redox sensing motif, which specifically recognizes cellooligomers with a degree of polymerization (DP) ≥ 3 and confers broad-spectrum disease resistance.

## Results

### Identification of a potential conserved redox sensor in LRR-Malectin RLKs

A comprehensive dataset of high-quality plant genomes was compiled from 1,177 plant species (**fig. S1A, Table S1**). Following rigorous manual curation, domain annotation, exclusion of erroneous results, and additional transcriptome searching for those genomes lacking detectable LRR-Malectin RLKs (**Table S2**), a total of 14,636 high-confidence LRR-Malectin RLKs were retained (**fig. S1B, Table S3**). Contrary to previous reports suggesting that the LRR-Malectin RLKs originated in *Physcomitrella patens* (*39*) or Chlorophyta (*40*), our analysis provides more robust evidence that this family of receptors was initially detected in *Spirogloea muscicola* (Zygnematophyceae) (**fig. S2**). Given that Zygnematophyceae is the sister lineage to embryophytes (land plants) (*41*), this supports the origin of LRR-Malectin RLKs in the common ancestor of Zygnematophyceae and embryophytes, potentially predating the colonization of land by plants. These RLKs could be divided into four well-supported clades (A-D) (**Fig. 1A,B**). Clades B/C are restricted to mosses and lycophytes with unknown functions. Clade D includes members from all major lineages of land plants, and can be further divided into three subclades of non-seed plants (D1-D3) and two subclades of seed plants (D4-D5). Interestingly, subclade D4 contains the orthologs of Arabidopsis IGPs and At1g56120 (**fig. S3**, hereafter termed as IGP-like 1, IGPL1), whereas subclade D5 contains the RFK members of Arabidopsis that have been reported to work synergistically in pollen-stigma interactions(**fig. S3**) (*5*). Although the functions of members in subclades D1/D2/D3 have not yet been reported, their relatively close phylogenetic relationships and similar gene structures suggest functional associations with subclades D4/D5 (**Fig. 1B, fig. S4A**). Above results indicate substantial potential functional diversification and a complex evolutionary history of LRR-Malectin RLKs.

**Fig. 1.**
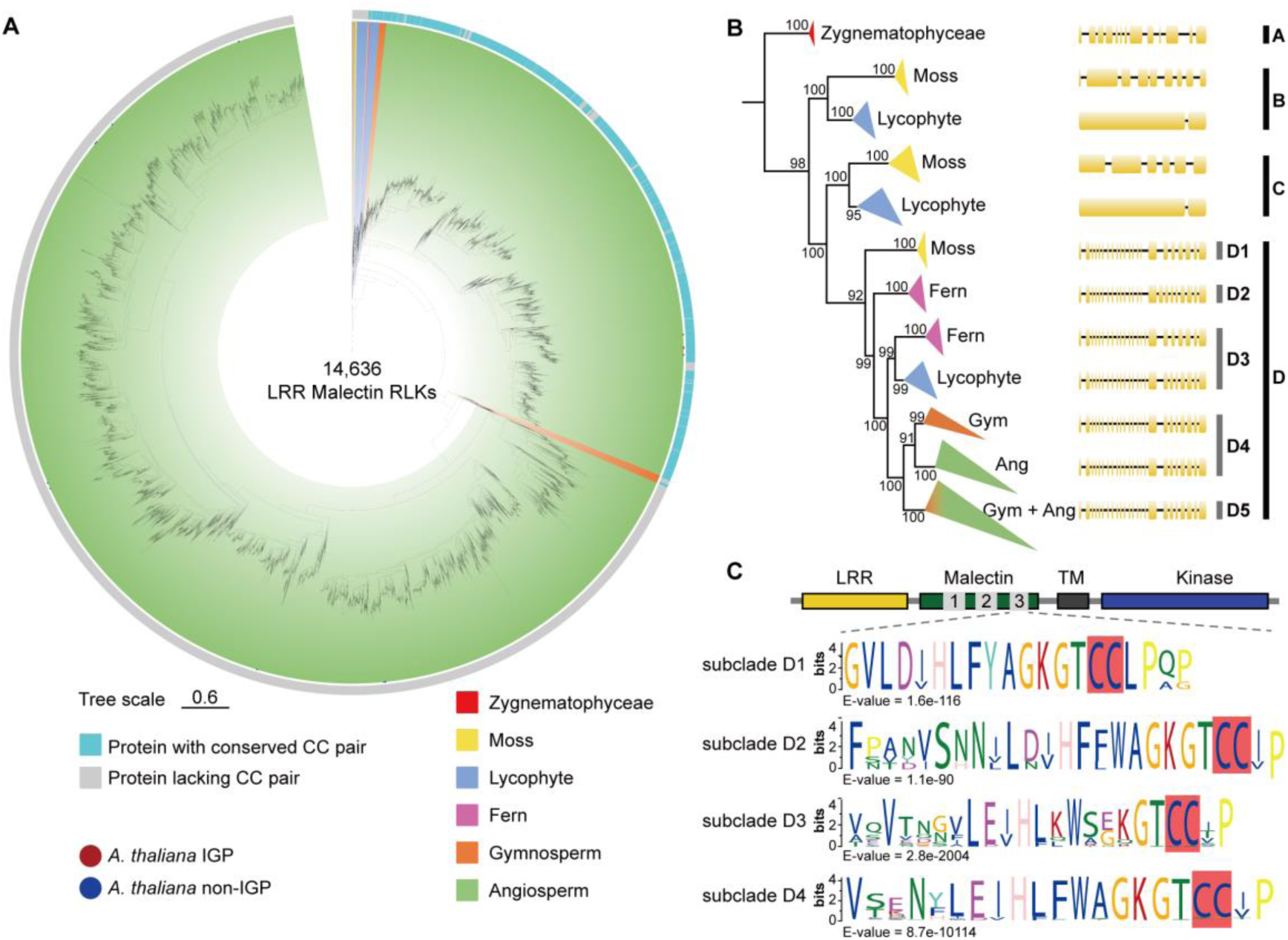
Identification of a potential conserved redox sensor in plant LRR-Malectin RLKs. (**A**) Phylogenetic tree of 14,636 LRR-malectin RLKs. The inner ring shows the topology. Algae, mosses, ferns, lycophytes, gymnosperms, and angiosperms are marked with different ground colors. IGPs and non-IGPs in Arabidopsis are annotated with red and blue circles, respectively, along the edge of the inner ring. (**B**) A simplified version of the phylogenetic tree, with schematic representations of gene structures attached next to each clade. (**C**) Conserved extracellular motifs of subclades D1/D2/D3/D4. The potential redox sensor CC pairs are marked with red background. LRR-malectin RLKs containing the CC pair or not are also indicated in the outer ring of phylogenetic tree in (**A**) by light blue or gray, respectively.

Since malectin was proposed as a ligand-binding domain, we next searched the conserved extracellular motifs in this structural domain of clades B-D to identify potential important residues in ligand perceptions. Notably, a motif containing a CC pair is highly conserved in subclades D1/D2/D3/D4, but is scarcely found in other clades. (**Fig. 1C, fig. S4B**). It is well known that the cysteine pair in the ectodomain (ECD) of RLKs primarily forms disulfide bonds to maintain structural stability. However, no disulfide bond is required for structural stability of MALECTIN (*24*) (**fig. S5A**). These data led us to hypothesize that this conserved CC pair in subclades D1/D2/D3/D4, like the two CC pairs (C421-C424 and C434-C436) in HPCA1 (*38*), may function as a potential redox sensor for extracellular ROS to regulate the function of receptors.

### Oxidation on C591/592 attenuates the function of IGP1

Next, we tested if the CC pair in IGP1 (C591/C592) from subclade D4 was a real redox sensor and whether redox-sensing alters the function of IGP1. We obtained T-DNA insertion mutants of IGPs and elucidated their role in plant disease resistance. Consistent with previous data (*14, 15*), cellotetraose (CEL4) induced significantly ROS burst and MAPK activity in Arabidopsis. These immune responses, however, were completely abolished in *igp1-2* and significantly reduced in *igp2/3-2* and *igp4-2* (**fig. S6A,B**). These genetic data showed that *IGP1* serves as the primary regulator of cellooligomer-induced immune signaling, with its homologs playing a contributory role in this pathway. In the *igp1-2* mutant, the expression of other *IGP* genes were significantly upregulated (**fig. S6C**), further suggesting functional redundancy among IGPs. As anticipated, all *igp*s mutants exhibited enhanced susceptibility to the virulent bacterium *Pseudomonas syringae* pv. *tomato* (*Pst*) DC3000 (**fig. S6D, E**), indicating that IGPs positively regulate disease resistance in Arabidopsis.

Notably, both CEL4 and H_2_O_2_ significantly upregulated *IGP*s gene expression in Arabidopsis (**Fig. 2A** and **fig. S7A**) and promoted the accumulation of IGP1 protein both in Arabidopsis and in *Nicotiana benthamiana* (**Fig. 2B,C** and **fig. S7B,C**). To examine whether H_2_O_2_ induces oxidative modification of IGP1, we transiently expressed and purified IGP1^ECD^-His and IGP1^ECD^-GFP from insect cell and *N. benthamiana*, respectively. MAL-PEG2000 (a 2 kD reagent that reacts with thiols of IGP1 in reduced state) assays demonstrated a dose-dependent increased oxidative modification of IGP1^ECD^-His upon prolonged H_2_O_2_ treatment (**Fig. 2D**). A simultaneous mutation of C591/C592 to Arg/Arg (CC-RR) abrogated the conjugation of MAL-PEG2000 with the IGP1^ECD^-His protein (**Fig. 2E**), demonstrating that this CC motif represents the primary site of free thiols *in vitro*. In contrast, under *in vivo* conditions, nearly all IGP1^ECD^-GFP was detected in an oxidized state in MAL-PEG2000 assays (**Fig. 2F**), suggesting that the CC motif is predominantly oxidized *in planta*. To further investigate whether higher oxidative modifications of IGP1 (S-sulfenylation/sulfinylation/sulfonylation/nitrosylation, excluding disulfide bond) occur in response to extracellular ROS elicited by CEL4, we purified IGP1 from overexpression lines of Arabidopsis treated with CEL4 and analyzed the protein by liquid chromatography tandem mass spectrometry (LC-MS/MS). The results showed that, among 10 extracellular Cys residues in IGP1, C591/C592 exhibited the highest-confidence oxidation signals with Cys S-sulfenylation/sulfinylation/sulfonylation/nitrosylation all identified (**Table S4**, **Fig. 2G** and **fig. S5B-E**), indicating that C591/C592 are also principal cysteines that sense H_2_O_2_ *in vivo*.

**Fig. 2.**
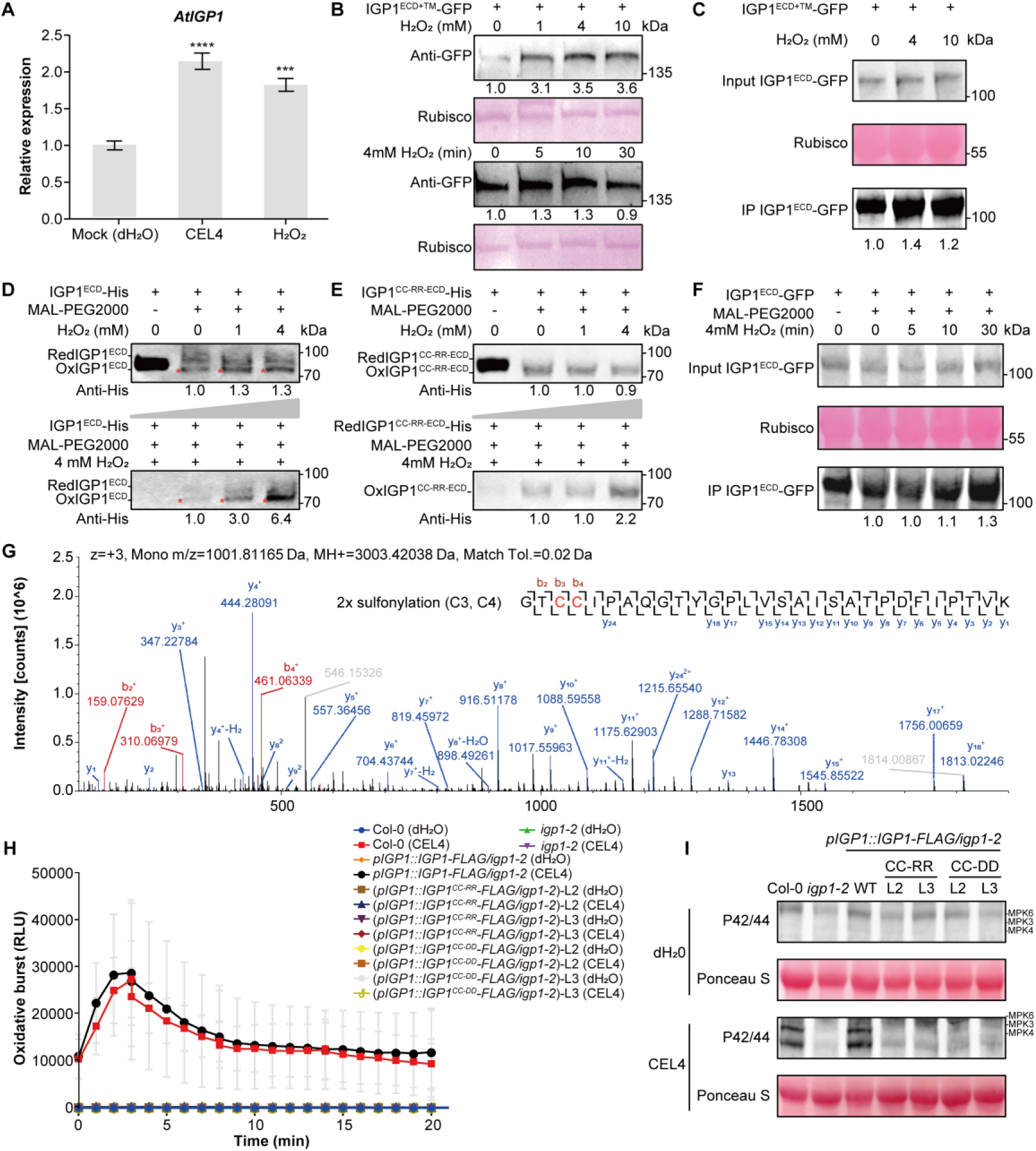
Oxidation on C591/C592 attenuates the function of IGP1. (**A**) CEL4 and H_2_O_2_ induce *AtIGP1* gene expression. qRT-PCR was performed to quantify the expression level of *AtIGP1* in Arabidopsis 40 min after treatment with 10 µM CEL4 or 20 µM H_2_O_2_ with dH_2_O as negative control. (**B**) Analysis of IGP1 protein expression in response to H_2_O_2_ treatment in Arabidopsis protoplasts. The expression level of the IGP1^ECD+TM^-GFP fusion protein was assessed in Arabidopsis protoplasts treated with varying concentrations of H_2_O_2_ (0, 1, 4, 10 mM) for 10 min, and with 4 mM H_2_O_2_ for different durations (0, 5, 10, 30 min). Protein loading was normalized using Ponceau S staining, with Rubisco serving as a loading control. The band intensity was quantified using ImageJ software. (**C**) Detection of IGP1 protein expression in *N. benthamiana* following H_2_O_2_ treatment. The IGP1^ECD^-GFP protein was transiently expressed in *N. benthamiana* via *Agrobacterium*-mediated infiltration. Leaves expressing the protein were treated with different concentrations of H_2_O_2_ (0, 4, 10 mM) for 30 min. The band intensity was quantified using ImageJ software. (**D** and **E**) *In vitro* analysis of the oxidation status of IGP1^ECD^ and IGP1^CC-RR-ECD^ proteins using the MAL-PEG2000 assay. The IGP1^ECD^-His and IGP1^CC-RR-ECD^-His proteins were expressed and purified from insect cells. For both proteins, samples were treated with different concentrations of H_2_O_2_ (upper panel) or samples with an increased concentration gradient were treated with 4 mM H_2_O_2_ (lower panel). The redox states of the proteins were detected by MAL-PEG2000 labeling, with Red and Ox indicating the reduced and oxidized states of the protein, respectively. The band intensity was quantified using ImageJ software. (**F**) *In vivo* analysis of the oxidation status of the IGP1 protein using the MAL-PEG2000 assay. The IGP1^ECD^-GFP protein was expressed in *N. benthamiana*. Leaves were treated with 4 mM H_2_O_2_ for different time periods (0, 5, 10, 30 min). The redox state of the IGP1 protein was then assessed by labeling with MAL-PEG2000. A control treatment without MAL-PEG2000 was included as a negative control. (**G**) LC– MS/MS spectrum recorded on the [M+3H]^3+^ ion at m/z 1001.81165 of the IGP1 CC peptide harboring two Cys-sulfonyl modifications. Predicted b- and y-type ions are listed. Detected ions, which indicate that C3 and C4 of the peptide are sulfonyl-modified, are labeled in the spectrum. Experiments were repeated three times with similar results. (**H** and **I**) The ROS burst (**H**) and MAPK activity (**I**) induced by 10 µM CEL4 were assessed in the complemented mutants CC-RR and CC-DD in the *igp1-2* mutant background. Two independent complementation lines were obtained for each mutant. Phosphorylation of MAPK was detected using an anti-p42/p44 antibody. Ponceau S staining of the blot was used to verify equal protein loading. Experiments were repeated three times with similar results.

To elucidate the function of the detected higher oxidative modifications of IGP1 C591/C592 in immunity, we introduced a C591D/C592D (CC-DD) mutation to mimic the S-sulfinylated state of IGP1 (**fig. S5E,F**) (*42*), along with the charge-reversed CC-RR mutant as a control. When expressed in the *igp1-2* mutant background, neither IGP1^CC-DD^ nor IGP1^CC-RR^ rescued the CEL4-induced immune response (**Fig. 2H,I**), indicating that higher oxidative modifications of the CC motif impede CEL4-triggered immunity.

### Cellooligomers with DP ≥ 3 activate IGP1

We next elucidated the mechanistic link between the redox-sensing and the receptor function of IGPs. The isothermal titration calorimetry (ITC) assays revealed that the insect cell-purified IGP1^ECD^ binds cellotriose (CEL3) and CEL4 with dissociation constants (K_d_) ∼ 1.61 and 1.48 μM, respectively. In contrast, no detectable interaction of IGP1^ECD^ with cellobiose (CEL2) (**Fig. 3A-C** and **fig. S8A**), explaining why little PTI response is induced by CEL2 (*16*). Notably, no binding was detected between IGP1^ECD^-tested hemi-celluloses (**Fig. 3C** and **fig. S8B, C**) or IGP4/IGPL1^ECD^-CEL4 (**fig. S8A**). Gel-filtration chromatography showed that IGP1^ECD^ and IGP1^ECD^-CEL4 eluted at a monomer position with molecular weight ∼85 kDa. Addition of CEL4 failed to induce IGP1^ECD^ heterodimerization with BAK1^LRR^ (**fig. S8D**). Furthermore, *in vivo* assays demonstrated that BAK1 was dispensable for CEL4-triggered ROS responses (**fig. S8E**). This finding aligns with previous reports that BAK1 was independent of cellooligomer-mediated PTI (*16, 43*)

**Fig. 3.**
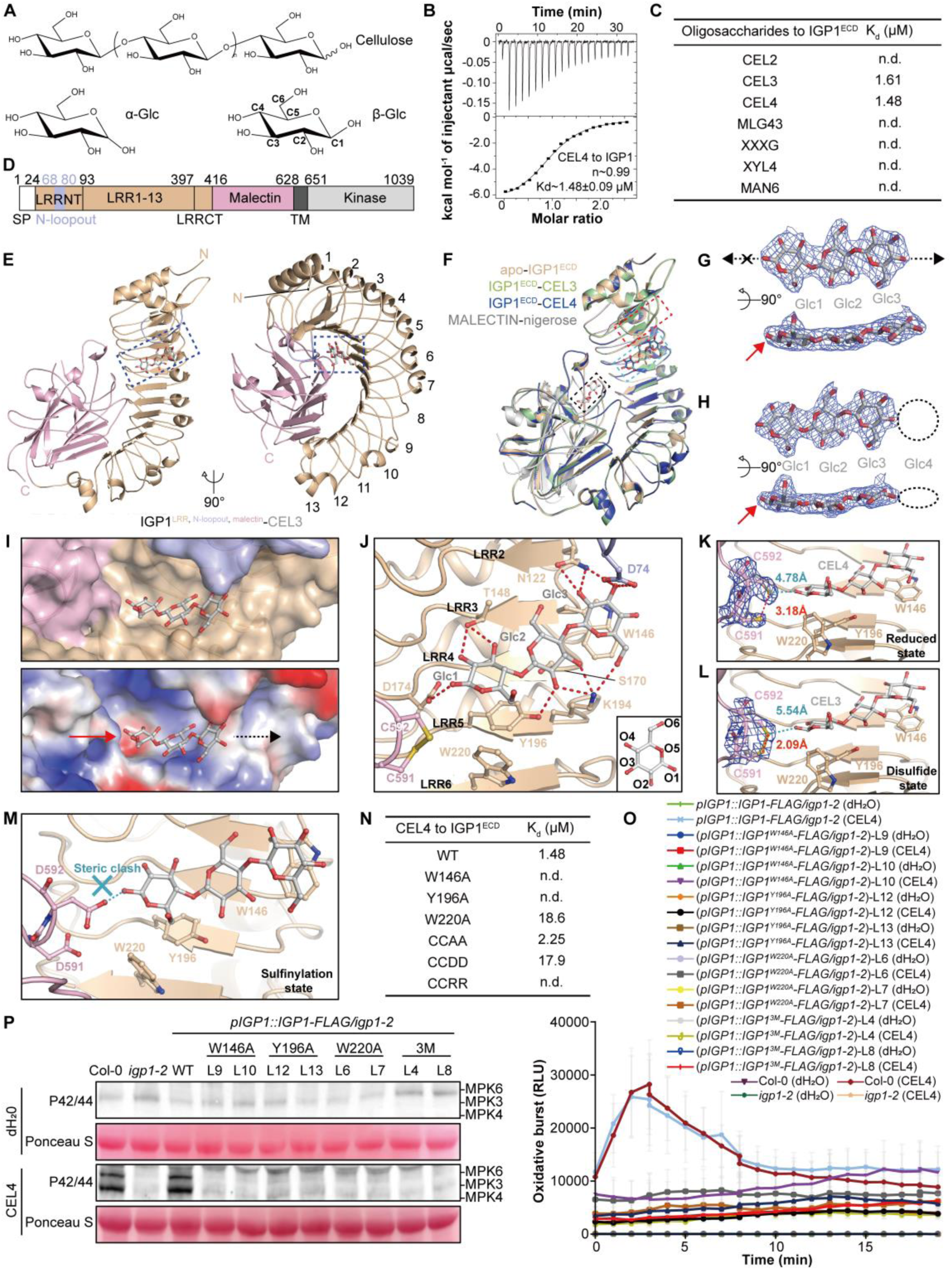
Mechanisms of cellooligomer recognition by IGP1. (**A**) Structural formula of cellooligomer and glucose isomers, α/β-glucose (Glc). The reducing end of cellooligomer adopting a changeable α/β conformation is depicted in wave line. C1-C6 in β-Glc are labelled as indicated. (**B**) Quantification of the IGP1^ECD^-CEL4 binding affinity at pH 6.0 by ITC. The binding constants (Kd values ± fitting errors) and stoichiometries (n) are indicated. (**C**) Summary of the affinity data on IGP1^ECD^-oligosaccharides interaction measured by ITC. Raw data are shown in Fig. 3B and fig**. S8A, C**. (**D**) IGP1 domain scheme: SP, signal peptide; LRR, leucine-rich repeat; LRRN/CT, LRR N/C-terminal cap; TM, transmembrane helix. (**E**) Cartoon diagrams of IGP1^ECD^-CEL3 complex in two orientations. The LRR1-13 of IGP1^ECD^ are numbered as indicated. CEL3 packing to LRR3-5 are framed in blue. N and C represent N and C termini, respectively. (**F**) Structural superposition of apo-IGP1^ECD^ with IGP1^ECD^– CEL3/4 and MALECTIN-nigerose (PDB: 2k46). The N-loopout motif of IGP1, CEL3/4, nigerose/predicted cellooligomer-binding pocket are framed in red, blue and black, respectively. (**G** and **H**) Electron density (2Fo-Fc, contoured at 1.5 sigma) around the cellooligomers in IGP1^ECD^-CEL3/4 structures shown in two directions. Red arrows indicate Glc1 adopting a β conformation. Black dash arrows indicate the permitted chain extension direction. Electron density corresponding to Glc4 in (**H**) was absent. (**I**) A cellooligomer binds to a half-open shallow surface groove on IGP1^ECD^. IGP1^ECD^ shown in surface and colored by domain scheme/electrostatics (upper/lower panels). White, blue and red indicate neutral, positive and negative electrostatics, respectively. (**J**) Details of the cellooligomer-IGP1^ECD^ interaction. Important residues for the interaction are shown. At the bottom right, O1-6 of Glc are indicated. In **G**-**J**, red/black arrows indicate the reducing end and the permitted extension direction of cellooligomer, respectively. (**K** and **L**) Electron density (2Fo-Fc) around C591-C592 of IGP1^ECD^-CEL3 (**K**) /CEL4 (**L**) contoured at 1.5 sigma. The atom distances between C591 and C592, and between C592 and the reducing end of Glc1 are labeled in red and blue, respectively. The disulfide/reduced states of C591-C592 are labeled. (**M**) Model on inhibition of cellooligomer perception by sulfinylated C592 (simulated by Asp) of IGP1. The cyan dash line indicates steric clash between sulfinylated C592 and the reducing end of cellooligomer. (**N**) Summary of the ITC data on IGP1^ECD^ WT/mutant-CEL4 interactions. Raw data are shown in Fig. 3B and fig**. S8F**. (**O** and **P**) The ROS burst (**O**) and MAPK activity (**P**) induced by 10 µM CEL4 were assessed in the complemented mutants W146A, Y196A, W220A, and the triple mutant 3M (W146A-Y196A-W220A). Two independent complementation lines were obtained for each mutant. Phosphorylation of MAPK was detected using an anti-p42/p44 antibody. Ponceau S staining of the blot was used to verify equal protein loading. In **B**, **O** and **P**, experiments were repeated three times with similar results.

The crystal structures of apo-IGP1^ECD^, apo-IGP4^ECD^, IGP1^ECD^-CEL3, IGP1^ECD^-CEL4 were resolved at resolutions of 2.49/2.65/2.74/2.69 Å, respectively (**Table S5**, **fig. S9A-C**). The IGP1^ECD^ structure harbors 13 tandem LRRs with a 12-amino-acid N-loopout embedding in LRRNT and a C-terminal malectin domain docked onto the LRR concave surface (**Fig. 3D, fig. S3B**). Structural superposition of IGP1^malectin^ with MALECTIN-nigerose (*24*) revealed a conserved core malectin architecture complemented by unique structure elements (η14, β27, β28) (**fig. S3B, S9D**). These elements along with a β36-β37 connecting loop mediate malectin-LRR interactions critical for overall structure stability (**fig. S3B, S9E**).

In the IGP1^ECD^-cellooligomer structures, IGP1^ECD^ and CEL3/4 form a 1:1 stoichiometry complex (**Fig. 3E, F**), corroborating the gel filtration data (**fig. S8D**). Structural comparison showed that CEL3/4 ligand-bound and apo-IGP1^ECD^ exhibited near-identical conformations (**Fig. 3F**), indicating no structural rearrangements upon CEL3/4 binding. The omit maps revealed that the reducing-end Glc1 of CEL3/4 adopts a β-conformation, while Glc1-3 maintain chair conformations in a slightly tilted plane (**Fig. 3G,H**). However, electron density corresponding to Glc4 was absent (**Fig. 3H**), indicating that Glc4 is flexible and not involved in interaction with IGP1^ECD^. These data explain the comparable IGP1^ECD^ binding affinities of CEL3/4 (**Fig. 3C**). Together, our structural and biochemical data establish CEL3 as the minimal unit required for IGP1 activation.

### Recognition mechanism of cellooligomers by IGPs

In contrast to the predictions of glycan-binding site in IGP1^malectin^ (**fig. S5A**), our structures unambiguously revealed that CEL3/4 occupy a previously unappreciated shallow groove composed of LRR3-5, N-loopout, and the malectin CC redox sensor (**Fig. 3I**), distal to the predicted sugar binding pocket (**Fig. 3F**). The N-loopout region has been shown to be a major structural determinant for ligand recognition of the RLPs RXEG1 (*44*), RPK2 (*45*) and certain LRR-RLPs (**fig. S10A**).

In the complex structures, Glc1-3 aligns parallel to β-strands in LRR3-5 through hydrophobic stacking and polar interactions. An aromatic triad (W220/Y196/W146, “WYW motif”) mediates pyranose ring stabilization via van der Waals contacts, dominating the cellooligomer perception (**Fig. 3J**). Complementary hydrogen bonding networks further reinforce binding: Glc1 O1/O2/O3 interact with D174 and T148; Glc2 O2/O3 form bonds with Y196 and K194; Glc3 O2/O3/O6 engage D74, N122, and K194 (**Fig. 3J**). Notably, D74 from the N-loopout and W146 creates a structural clamp that stabilizes Glc3. No oxidizing or reducing agents were added to the crystallization buffer, the CC motif in CEL4-bound IGP1^ECD^ was observed in a reduced state (**Fig. 3K**). This contrasts with the CC motifs adopting a disulfide state in CEL3-bound IGP1^ECD^, apo-IGP1^ECD^ and apo-IGP14^ECD^ as indicated by electron density and the distance between the two cysteine residues in the CC motif (**Fig. 3L** and **fig. S9F, G**). The CC motif forms no direct contact with CEL3/4. However, higher oxidative modification of the CC motif would increase the sizes of the two cysteine residues thus creates steric clash with cellooligomers as shown by modeling study (**Fig. 3M**).

Although the IGP1-4/L1 shares a conserved WYW-CC motif and a similar ligand-binding pocket (**fig. S9H**), the cellooligomers binding residues of IGP1 are not conserved in IGP2-4/L1. Notably, the critical cellooligomer binding residues such as D74, A172, D174, S170, K194 found in IGP1 have been substituted with other residues (**fig. S3B**). This divergence likely underlies the mechanism of cellooligomers specific recognition by IGP1.

Structural comparison revealed the mechanisms of specific recognition of cellulose and chitin by their respective receptors. Although chitin shares a cellulose backbone (but carries a modification on C2) (**fig. S10B**), structural superposition of IGP1-CEL3 with OsCEBiP-NAG3 (*46*) showed that IGP1 recognizes CEL3 from an inverted orientation relative to NAG3 (**fig. S10C**). In further contrast to the chitin receptors such as LysM receptors (*46–48*) that permit bidirectional extension of bound chitooligosaccharides (**fig. S10D**), IGP1 interacts with the reducing end of cellooligomers, thus sterically blocking cellooligomer chain extension at this site (**Fig. 3I**). These structural differences explain the requirement of the intact reducing end for cellooligomer perception. Molecular modeling confirmed that Berberine bridge enzyme-catalyzed Glc1 ring oxidation opening creates steric incompatibilities with T148, D174, and W220 (**fig. S10E**), consistent with biochemical evidence demonstrating loss of elicitor activity of oxidized cellooligomers (*49, 50*).

The molecular basis for ligand specificity was further elucidated through systematic structural comparisons. Key exclusion mechanisms prevent recognition of related hemi-cellulose DAMPs: in XXXG (a hydrolysate of xyloglucan carrying a CEL4 backbone with O6 xylosylation), the critical O6 position of Glc2 faces inward within the binding pocket—xylose substitution induces steric clashes with N122. By comparison, the lack of C6-O6 would prevent xylotetraose (XYL4) from interactions of Glc1/3 with W220/K194, respectively. The stereochemical inversion at O2 in mannohexaose (MAN6, C2 epimer of CEL6) disrupts the precisely positioned hydrogen bonds of Glc1/2/3 with T148/Y196/N122, respectively (**Fig. 3J**). Collectively, these findings reveal the mechanism that enables plants to distinguish intact cellooligomers from structurally similar polysaccharides.

### Mutagenesis of the IGP1-cellooligomer complex

To verify our structures, we introduced alanine substitutions into the WYW-CC motif in IGP1, and evaluated their effects on the IGP1-CEL4 interaction using ITC. The ITC data showed that the W220A mutation reduced the binding affinity, yielding a K_d_ of ∼18.6 μM and no interaction was detected for the W146A or Y196A variants (**Fig. 3N** and **fig. S8F**). To further support our structural data, we made transgenic Arabidopsis plants expressing IGP1-FLAG driven by its native promotor in *igp1-2*. As shown in **Fig. 3O,P**, the IGP1^WT^ transgenic plants fully recovered CEL4-triggered ROS burst and MAPK activation to WT levels. In contrast, the IGP1^W146A^, IGP1^Y196A^, IGP1^W220A^ and IGP1^3M^ (Y196A/W146A/W220A) transgenic plants failed to restore these immune responses (**Fig. 3O,P**), underscoring the essential role of these residues in IGP1-triggered immunity. Similarly, ITC analyses revealed that IGP1^ECD, CC-DD^ interacted with CEL4 with a increased K_d_ of ∼17.9 μM and the CC-RR variant exhibited no detectable binding. These results align with the inability of IGP1^CC-RR^ and IGP1^CC-DD^ to restore CEL4-induced immunity in planta (**Fig. 2H,I**). Substitution of the CC motif with AA would not substantially impact the CEL4-binding activity of IGP1, because this mutation results in no steric clash with CEL4. Indeed, IGP1^ECD, CC-AA^ displayed a K_d_ of ∼2.25 μM, similar to that of wild-type IGP1 ^ECD^ (**Fig. 3N**).

Collectively, these biochemical and functional evidence validates the structural model of the IGP1–cellooligomer complexes and supports an essential role of their specific interaction in cellooligomer-induced immune signaling.

### Origin and evolution of IGPs: structural conservation and divergence

IGP1-CEL structure-based sequence alignment confirmed that nearly all members in subclades D1/D2/D3/D4 are IGP orthologs containing a putative cellooligomer-binding pocket (**fig. S11A**). Ultimately, a total of 4,558 IGPs were identified from 1,037 species (**Table S1,S3**). The earliest occurrence of IGPs were detected in the moss *Ceratodon purpureus*, indicating that they likely originated in the common ancestor of embryophytes and arose during plant terrestrialization (**Fig. 4A** and **fig. S2**). The origin of IGPs may be linked to elevated pathogen pressure in terrestrial environments. Consistent with recent studies of plant PRR repertoires (*51, 52*), further analysis of angiosperm IGPs suggests that the evolution of IGPs may have also been shaped by a combination of genomic variation (**fig. S12A**), lineage-specific expansions and contractions (**fig. S12B,C**), and environmental selection pressures and trophic differentiation (**fig. S13A-C**).

**Fig. 4.**
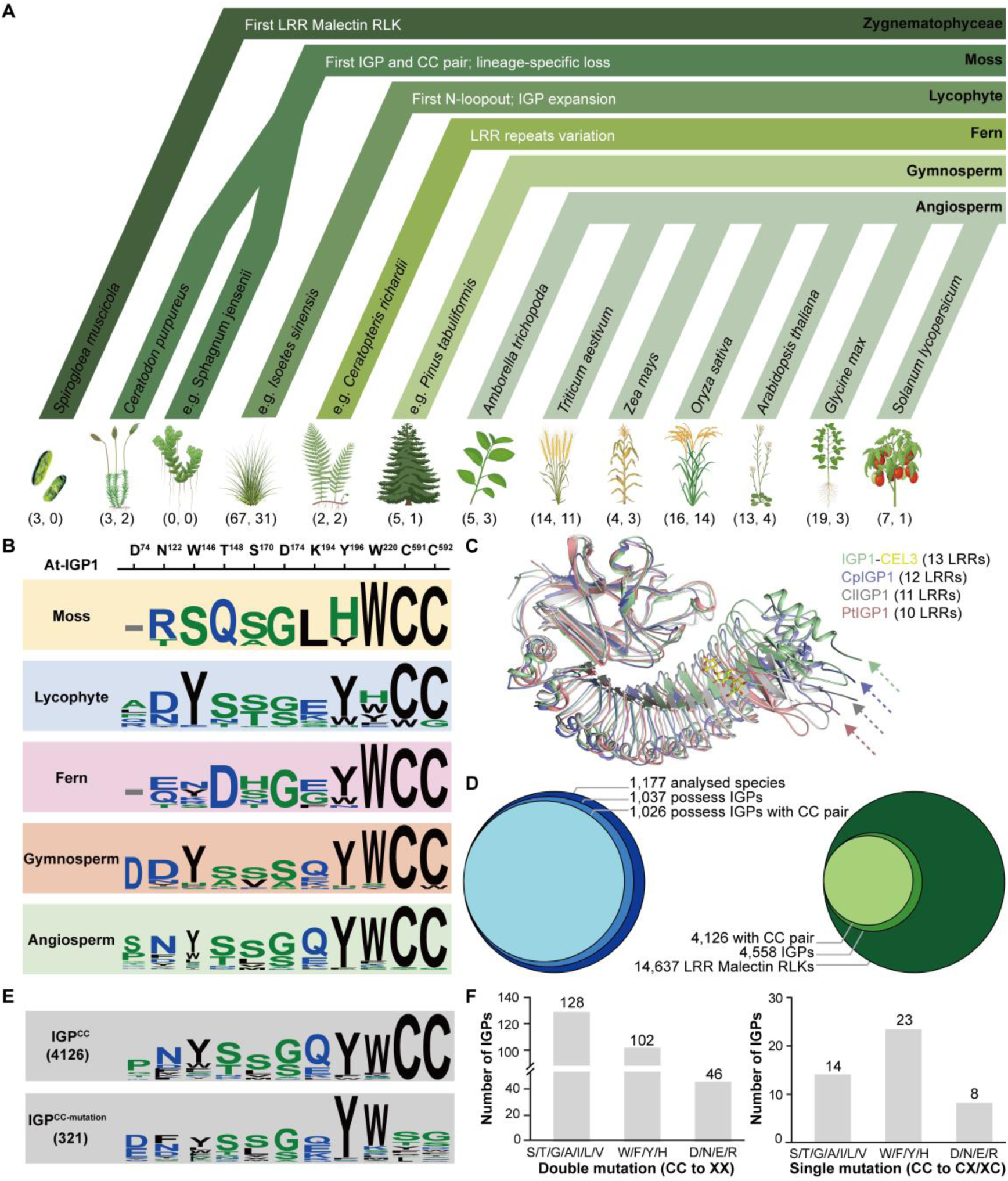
Structural conservation and divergence in IGP evolution. (**A**) Evolutionary framework of IGPs. The major evolutionary events are indicated in the respective plant groups. Different plant groups are visually distinguished by applying distinct background colors. The numbers of LRR-malectin RLKs and IGPs from selected representative species are annotated in brackets. (**B**) Amino acid usage frequencies at the cellooligomer-binding pocket across different plant groups. The corresponding positions in IGP1 are indicated above each site. The absence of the N-loopout motif is indicated short grey dash. W/Y/F/H can be considered functionally equivalent for binding cellooligomer. (**C**). Structural superposition of IGP1-CEL3 with CpIGP1/ClIGP1/PtIGP1. Cp: *Ceratodon purpureus*; Cl: *Cunninghamia lanceolata*; Pt: *Pinus tabuliformis*. The arrows indicate the first LRR of each IGP. (**D**) Distribution numbers of IGP1 and IGP1 with the CC motif across analyzed species and LRR-malectin RLK family. (**E**) Amino acid usage frequencies at the cellooligomer-binding pocket across IGPs with or without the CC motif. (**F**) Statistics of 321 structurally intact IGPs with natural mutations in the CC motif. Left: complete mutation of the CC motif including both two Cys mutated to S/T/G/A/I/L/V, at least one Cys mutated to W/Y/F/H, and mutations containing D/N/E/R but not W/Y/F/H. Right: a single Cys of the CC motif mutated to S/T/G/A/I/L/V, W/Y/F/H, and D/N/E/R.

IGP orthologs exhibit a high degree of structural conservation for cellooligomer perception: analysis of amino acid usage frequency within the putative cellooligomer-binding site indicates the core ‘WYWCC’ motif identified in Arabidopsis is conserved in lycophytes and seed plants, while a ‘YWCC’ motif is conserved in mosses and ferns (**Fig. 4B** and **fig. S11A, S14A,B**). Notably, IGP^LRR^ structures exhibit significant lineage-specific divergence, primarily manifested as variations in LRR number and acquisition of the N-loopout motif. Mosses IGPs typically contain 12 LRRs and lack the N-loopout motif. Ferns similarly lack the N-loopout motif, but exhibit either 12 or 13 LRRs. In contrast, lycophytes and seed plants predominantly exhibit IGPs with 13 LRRs and the N-loopout motif (**fig. S11A, S14A** and **Table S6**). In the meantime, rare IGP orthologs with 10 or 11 LRRs were identified (**Table S6**). Structural alignment of IGPs with 10-13 LRRs shows that major difference occurs at the N-terminal LRRs surrounding cellooligomer-binding pocket (**Fig. 4C** and **fig. S11B**), potentially reflecting selective adaptation for recognizing oligosaccharides of varying sizes. Furthermore, since IGP1 CC motif substitution affects IGP1-cellooligomer interaction (**Fig. 3N**), we investigated the natural mutation landscape of the CC motif within IGPs. Of the 432 IGPs exhibiting variations at CC motif, 321 with intact structures were selected for subsequent analyses (**Fig. 4D,E**). Complete variation at the CC motif was observed in 276 IGPs. (**Fig. 4F**). Among these fully mutated motifs, approximately 46.3% were both mutated to residues S/T/G/A/I/L/V that abolish redox sensing capability while preserving oligosaccharide recognition with a high probability. Approximately 37.0% contained at least one mutation to residues W/F/Y/H, potentially enhancing oligosaccharide sensing. The rest mutations containing like D/N/E/R, which impair oligosaccharide sensing, accounted for only approximately 16.7%. Furthermore, there are 45 IGPs with a single cysteine residue mutated; the distribution of mutation types among these IGPs mirrored that observed in the fully mutated group (**Fig. 4F** and **Table S7**). Collectively, these findings strongly suggest the importance of preserving the redox-sensing capacity in IGP evolution and that natural mutations within the IGP CC motif are constrained by rigorous selection.

### IGP-cellooligomer immune modules confer broad-spectrum disease resistance

We next determined whether the identified conserved IGP orthologs in crop species confer disease resistance. A genome-wide association study (GWAS) was conducted on 349 Chinese wheat (*Triticum aestivum*) accessions to identify IGP loci associated with Fusarium crown rot (FCR) and leaf rust (LR) resistance, using the Wheat 660K SNP array. We identified that SNP locus in the gene *TraesCS7A03G0083800* (designated as *TaIGP1*), which belongs to the *TaIGP* family (**fig. S15**), was significantly associated with FCR resistance across three environments and with LR resistance in one environment, suggesting its potential role in multi-disease resistance in wheat (**Fig 5A**). We therefore investigated whether *TaIGP1* also confers resistance to powdery mildew (Pm), another economically important wheat disease. We silenced *TaIGP1* expression via virus-induced gene silencing (VIGS) in the moderately Pm-resistant cultivar Zhengmai9023. qRT-PCR confirmed successful *TaIGP1* knockdown 14 days post-barley stripe mosaic virus (BSMV) inoculation. Upon infection with powdery mildew isolate E09, the silenced plants displayed a significantly higher disease index than controls, demonstrating that *TaIGP1* is a positive regulator of wheat powdery mildew resistance. (**Fig 5B**).

**Fig. 5.**
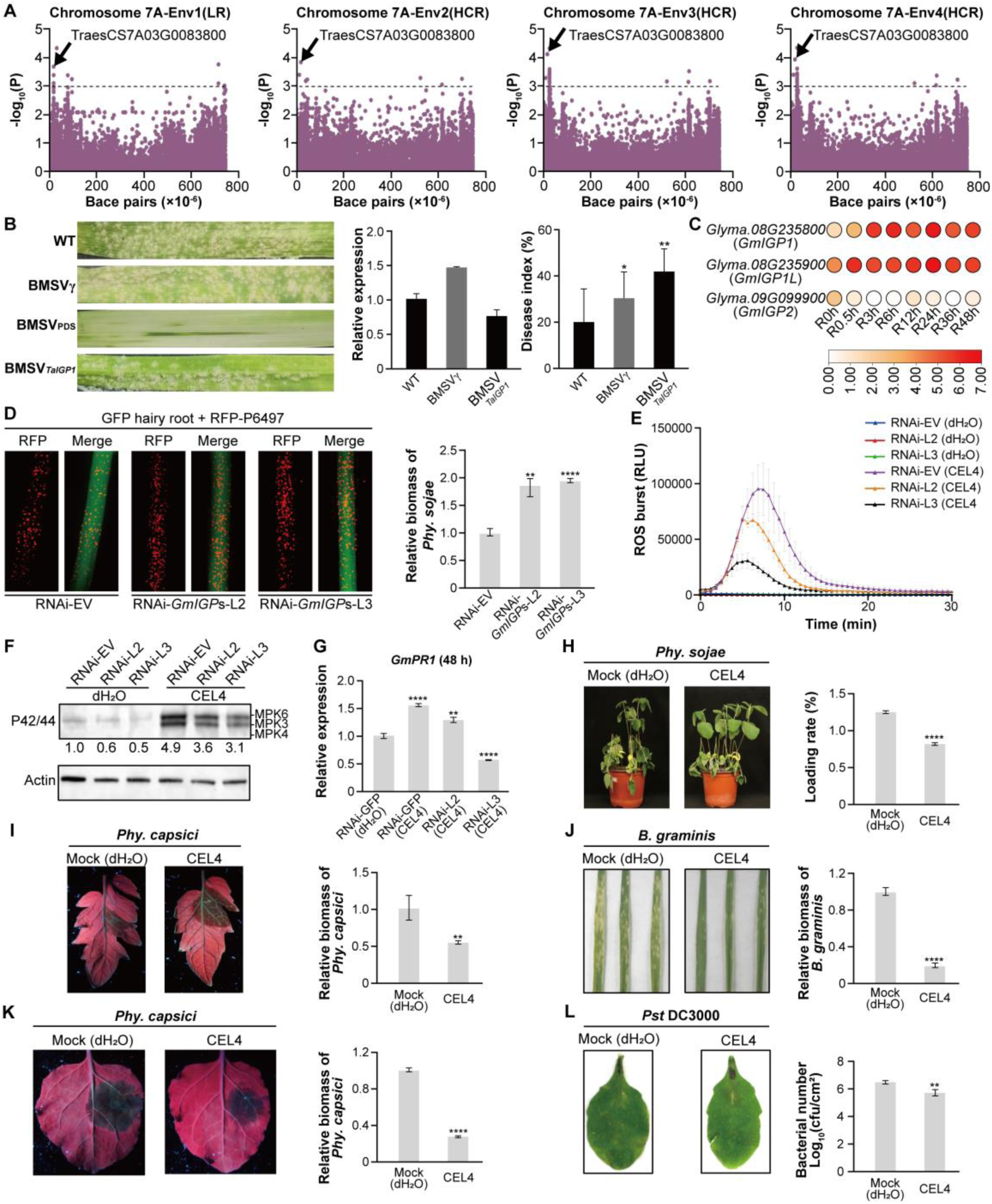
IGP-cellooligomer immune modules confer broad-spectrum disease resistance. (**A**). A genome-wide association study (GWAS) identified *TaIGP1* as a locus associated with FCR and LR resistance. The Manhattan plot shows that significant SNP within *TaIGP1* (arrow) was located on chromosome 7A and was consistently detected across four environments. (**B**). *TaIGP1* silencing compromises powdery mildew (Pm) resistance in wheat. Virus-induced gene silencing (VIGS) was performed in the moderately Pm-resistant wheat cultivar Zhengmai9023. The Pm disease phenotype, relative expression levels of *TaIGP1*, and the disease index are shown for wild-type (WT) plants, control plants inoculated with the empty BSMV vector (BSMV_γ_), and *TaIGP1*-silenced plants (BSMV*_TaIGP1_*). (**C**) Transcriptional patterns of *GmIGP1*, *GmIGP1L*, and *GmIGP2* genes in soybean during different stages of infection. The color intensity represents the transcript level, with darker red indicating higher transcriptional activity. Transcript expression levels were normalized using log2 transformation. (**D**) Co-silenced *GmIGP*s in soybean hairy roots reduce soybean resistance against *Phy. sojae*. Transgenic hairy roots expressing *GFP* (RNAi-EV), co-silenced *GmIGP1*, *GmIGP1L*, and *GmIGP2* (RNAi-*GmIGP1*/*IGP1L*/*IGP2*-Line 2 and RNAi-*GmIGP1*/*IGP1L*/*IGP2*-Line 3) constructs (GFP control or genes RNAi) were inoculated with *Phy. sojae* zoospores expression red fluorescence protein (RFP) (left). Oospore production 48 hpi is shown. Six independent experiments exhibit similar results. Left, RFP; right, merge (RFP + GFP). Scale bars, 0.2 mm. The relative biomass of *Phy. sojae* in infected hairy roots, as measured by genomic DNA qPCR and normalized to the GFP control (right). (**E**) ROS production triggered by CEL4 (10 μM) in hairy roots of soybean plants co-silenced *GmIGP*s with dH_2_O as negative control. Mean RLU (Relative Luminescence Unit) (± SD) is shown (n=4). (**F**) CEL4 triggered MAPK phosphorylation in co-silenced *GmIGP*s transgenic soybean hairy roots at 10 min with dH_2_O as negative control. Total protein was analyzed by immunoblot with an antibody for phosphorylated MPK6/3/4 (P42/44). Actin was used as a loading control. The total band intensities were quantified using Image J. (**G**) qRT-PCR analysis of *GmPR1* gene expression induced by CEL4 in co-silenced *GmIGP*s transgenic soybean hairy roots at 48 h with dH_2_O as negative control. (**H**) Visual symptoms 4 dpi with *Phy. sojae* P6497 on the stem of soybean plants, which were pretreated separately with mock (dH_2_O) and CEL4 (10 μM). Representative photographs are shown (left). The lodging rate of soybean plants was statistically analyzed (right). (**I**) Visual symptoms 2 dpi with *Phy. capsici* on tomato leaves, which were pretreated separately with mock (dH_2_O) and CEL4 (10 μM). Representative photographs are shown (left). Relative biomass of *Phy. capsici* infecting tomato leaves (at 2 dpi) as measured by genomic DNA quantitative PCR, and normalized to *Phy. capsici* (dH_2_O) (right). (**J**) Visual symptoms 2 dpi with *B. graminis* on wheat leaves, which were pretreated separately with mock (dH_2_O) and CEL4 (10 μM). Representative photographs are shown (left). Relative biomass of *B. graminis* infecting wheat leaves (at 2 dpi) as measured by genomic DNA quantitative PCR, and normalized to *B. graminis* (dH_2_O) (right). (**K**) Visual symptoms 2 dpi with *Phy. capsici* on *N. benthamiana* leaves, which were pretreated separately with mock (dH_2_O) and CEL4 (10 μM). Representative photographs are shown (left). Relative biomass of *Phy. capsici* infecting *N. benthamiana* leaves (at 2 dpi) as measured by genomic DNA quantitative PCR, and normalized to *Phy. capsici* (dH_2_O) (right). (**L**) Visual symptoms 2 dpi with *Pst* DC3000 at 5×10^5^ cfu/mL on Arabidopsis Col-0 leaves, which were pretreated separately with mock (dH_2_O) and CEL4 (10 μM). Representative photographs are shown (left). Bacterial growth is shown as mean ± SD (*n* = 6).

In parallel, we identified three *GmIGP* homologs in soybean (*Glycine max*)— *Glyma.08G235800* (*GmIGP1*), *Glyma.08G235900* (*GmIGP1L*), and *Glyma.09G099900* (*GmIGP2*) (**fig. S16A**). Transcriptome analysis of *Phytophthora sojae*-infected soybean revealed that *GmIGP1* and *GmIGP1L* were significantly upregulated, whereas *GmIGP2* expression was undetectable (**Fig 5C**). To probe their functional role in cellooligomer-triggered immunity, we co-silenced all three *GmIGP*s in soybean root hairs. Two independent silencing lines (L2 and L3) exhibited approximately 50% reduction in transcript levels for all three genes (**fig. S16B**). Silencing of *GmIGP*s significantly enhanced *Phy. sojae* infection (**Fig 5D**), indicating that *GmIGP*s positively regulate soybean resistance against *Phy. sojae*. Furthermore, CEL4-induced ROS burst, MAPK activity, and expression of the defense-related gene *GmPR1* were significantly reduced in *GmIGP*s-silenced root hairs compared to the RNAi empty vector control (**Fig 5E-G**), demonstrating that *GmIGP*s are essential for cellooligomer-triggered immune signaling.

We next investigated whether cellooligomers confer broad-spectrum disease resistance across diverse plant species. Pretreatment with CEL4 effectively enhanced soybean resistance against *Phy. sojae*, *Fusarium graminearum* and *Phakopsora pachyrhizi* (**Fig. 5H** and **fig. S16C-E**). Similarly, CEL4 application induced significantly induced disease resistance in tomato against *Phytophthora capsici*, wheat against *Blumeria graminis f. sp. Tritic*, *N. benthamiana* against *Phy. capsici*, and Arabidopsis against *Pst* DC3000 (**Fig 5I-L**). Together, these findings highlight the potential of cellooligomers as novel biopesticides and highlight crop IGPs as promising targets for engineering broad-spectrum disease resistance.

## Discussion

Plant RLKs harbor diverse oligosaccharide-binding domains, including LysM(*53*), L-lectin(*54*), G-lectin(*55*), malectin(*24*), DUF26(*56*), etc. Unexpectedly, our study unveils that IGP1^LRR^ functions as the platform for cellooligomer recognition through a conserved WYW motif. This finding, coupled with a recent preprint report identifying LRR-RLK ARMs as receptors for rhamnogalacturonan-I (*57*), provides compelling evidence establishing LRR domains as novel platforms for oligosaccharide recognition. Furthermore, we demonstrate that the CC motif of IGP1^malectin^ functions as a redox-sensor that directly regulates IGP1-cellooligomer interaction. Multiple lines of evidence indicate that oxidation of the two cysteine residues within this motif decreases cellooligomer binding via a steric hindrance mechanism. Notably, oxidized cellooligomers were found to lose their elicitor activity, indicating that oxidative modifications impair both ligand functionality and receptor-ligand interaction in the IGP1-cellooligomers pathway. Collectively, these findings highlight a dual regulatory role for oxidation in dampening the IGP1-mediated cellooligomer signaling pathway. This redox-dependent modulation may serve as a crucial feedback mechanism to prevent overactivation of plant immunity, thereby maintaining cellular homeostasis under stress conditions (**fig. S17**).

During the transition to a terrestrial environment, plants encountered a dramatic increase in both abundance and diversity of pathogens, which likely drove the innovation and evolution of cell wall oligosaccharide receptors like IGPs. Given that cellooligomers function not only as immune elicitors but also as inherent products of cell wall remodeling during normal growth and development, the variation in IGP orthologs number or LRR structure across plant lineages likely stems from more complex evolutionary drivers. The expansion and structural change of IGP families within specific plant lineages may signify the acquisition of unique oligosaccharide recognition capabilities. This concept is exemplified by EPS3, a *Lotus japonicus* LysM-RLK, which represents a unique class of plant oligosaccharide receptors specifically recognizing bacterial exopolysaccharides yet unresponsive to chitooligosaccharides (*58, 59*). Therefore, elucidating the adaptive evolutionary mechanisms underpinning the IGPs functional diversification across plant taxa also warrants further investigation.

Distinct from the NAG-CERK interaction, our study identified a novel unidirectional binding mechanism of cellooligomer perception by IGPs. Intriguingly, given the identical backbone structure of chitooligosaccharides and cellooligomers, and the conservation of both IGP and CERK receptors across higher plants, potential crosstalk between these signaling pathways warrants future investigations. Supporting this notion, interactions between OsCERK1 and CEL3/4 have been detected (*60*). In this regard, investigating whether IGPs participate in chitooligosaccharide-triggered immunity is highly pertinent. Additionally, it would be insightful to determine if LysM receptors can function as primary cellooligomer receptors in those plant lineages that have specifically lost IGP orthologs.

Ligand-induced heterodimerization of RLKs/RLPs with co-receptor such as SERKs represents a fundamental mechanism for most identified LRR-RLK/RLPs (*61*). SERKs have been reported to act as co-receptors for RKF1 and RKFL1/2/3 (*6*), but not for IGP1 based on our *in vitro* and *in vivo* data (*16, 43*). This raises a key question: how do cellooligomers induce IGP1 homodimerization or heterodimerization with IGP2-4/L1. Since our phylogenetic analyses reveal that IGP and RKF orthologs constitute two distinct functional subclades, it would therefore be valuable to investigate the structural differences underlying these divergent receptor activation mechanisms in the future.

The conventional agrochemicals are often costly and pose significant risks to human health and the environment. The eco-friendly plant- and fungal-derived cell wall oligosaccharides act as effective plant immunostimulants, holding immense promise as next generation of biopesticides. This potential is already being realized commercially: oligosaccharide-based products such as FytoSave^®^, Planticine^®^, Pectimorf^®^ (OGs), FytoSol^®^ (OGs and chitosan oligomers), and Vacciplant^®^ (β-1,3-glucans) are successfully deployed, demonstrating enhanced broad-spectrum resistance against diverse phytopathogens across multiple plant species (*62–66*). Our study provides foundational evidence supporting this trend: cellooligomers trigger PTI responses and bolster broad-spectrum resistance through a conserved mechanism in model plants spanning key crop families (Poaceae, Brassicaceae, Leguminosae and Solanaceae) via IGPs. We believe these findings establish a robust scientific basis and pave the way for the accelerated development and optimization of biopesticides based on novel plant cell wall oligosaccharides.

## Acknowledgments

We thank the staff of Beamline BL02U1, BL10U2, BL18U1 and BL19U1 at Shanghai Synchrotron Radiation Facility (SSRF) and X-ray crystallography platform, National Protein Science Facility, Tsinghua University, for assistance in X-ray diffraction data collection. We also thank the Mass Spectrometry and Metabolomics Core Facility of the Biomedical Research Core Facility, Westlake University, for the support in LC–MS data collection. We are grateful to Prof. Jijie Chai (Westlake University) for his guidance throughout this research project and Prof. Xing Liu (Wuhan University) for kindly providing access to the unpublished genome data of *Isoetes yunguiensis*.

## Author contributions

Y.X., X.W., Y.W., A.J., and G.S. conceived and conceptualized the study and designed the experiments. Y.X. A.J., and J.D. performed protein expression, protein purification and ITC assays. Y.X. performed crystallization screening. Y.X and Z.H. performed X-ray and structural data analysis. P.W. and X.W. performed phylogenetic analysis. G.S., J.H., X.M., X.L., and Q.Z. conducted the assessment of protein oxidative modifications and plant immune responses. G.S., J.H., and X.M. were responsible for the Arabidopsis transformation. X.M., X.L., and Q.Z. performed the soybean root hair transformations. N.X., and Z.Z. performed the analysis of the transcriptome data from *Phy. sojae*-infected soybean plants. M.F. performed the collection and analysis of MS data. G.S., A.J., J.D., X.M, X.L., L.G., Y.Y., M.F. and M.C. performed broad-spectrum disease resistance assays. Y.X., G.S., P.W. and A.J. wrote the manuscript with input from all authors.

## Data and materials availability

All data are available in the manuscript or the supplementary materials, or the listed Protein Data Bank (PDB) files. The atomic coordinates and structure factors have been deposited in the Research Collaboratory for Structural Bioinformatics PDB. The PDB codes for the crystal structures of apo-IGP1^ECD^, apo-IGP4^ECD^, IGP1^ECD^-CEL3 and IGP1^ECD^-CEL4 are 9WT4, 9WT5, 9WT6 and 9WT7, respectively. Genomic data were downloaded from publicly available databases, including NCBI, Phytozome, Ensembl, and NGDC. Detailed information is provided in the Supplementary Materials.

## Materials and Methods

### Identification of LRR-malectin RLKs

We first identified all LRR-malectin RLKs and then selected IGPs based on phylogenetic inference. To investigate the origin and evolution of IGP, we collected 1,177 genomic datasets spanning algae, mosses, ferns, lycophytes, gymnosperms, and angiosperms. For ease of reference, we retained the original protein names and added the corresponding species names as a prefix, formatted as ‘species--proteinID’. A total of 46,929,358 primary gene models of proteins were filtered and used for subsequent analyses. To ensure the accuracy of protein identification, we established a robust and comprehensive screening pipeline (**fig. S1B**). The previously published identification methods of LRR RLKs (*40, 51, 52, 67*) was optimized and applied in this study. Firstly, the HMMER 3.3.2 software was employed to detect the presence of the kinase domain (*68*) (Pfam: PF00069 or PF07714), the malectin domain (Pfam: PF11721), and the LRR domain (Pfam: PF00560, PF01462, PF01463, PF01816, PF07723, PF07725, PF08263, PF12799, PF13306, PF13516, PF13855, PF14580, PF18805, PF18831, PF18837, PF23286, or PF23598). After taking the intersection of three lists above, 18,075 proteins passed the HMMER search. We then used InterProScan 5.69 software to further annotate the detailed domains of these proteins, with the following parameters: -dp -f GFF3 -iprlookup -pa -appl CDD, COILS, Gene3D, HAMAP, MobiDBLite, PANTHER, Pfam, PRINTS, PROSITEPATTERNS, PROSITEPROFILES, SFLD, SMART, SUPERFAMILY, TIGRFAM, tmhmm, signalp_euk, phobius (*69*). This step enabled the precise exclusion of proteins with multiple kinases or malectin domains, non-LRR-malectin RLK domains, or incorrectly ordered domains. 14,481 proteins were filtered out, and each was manually examined to ensure accuracy. For species in which no LRR-malectin RLK was identified from the available genome data, we additionally downloaded their transcriptome datasets and conducted a *de novo* assembly to perform cross-validation (**Table S2**). Following de-redundancy and gene annotation to obtain the protein sequences, the remaining identification steps were performed as described above. Finally, a total of 14,636 proteins were identified as high-confidence LRR-malectin RLKs. Protein structures of selected candidates were predicted by AlphaFold 3(*70*).

### Taxonomic information and phylogenetic tree of species

The taxonomic information of species was obtained from NCBI. After identifying all LRR-malectin RLKs, we found that it first emerged in Streptophyta. Therefore, more ancient species were excluded from the phylogenetic analysis. The *Volvox reticuliferus*, which is belonging to Chlorophyta, was set as outgroup. Batch reciprocal blast method was employed to identify orthologs from different species (*71*). Iqtree 2.4.0 software was used to construct phylogenetic tree based on 67 single copy orthologs, with the parameters of -m MFP --mset LG --rclusterf 10 -B 1000 --alrt 1000 (*72*). Phylogenetic relationships among angiosperm species were generated with the V.PhyloMaker2 package (*73*). The cartoon images of plant species were obtained from online websites https://app.biorender.com/.

### Phylogenetic analysis of LRR-malectin RLKs

A total of 14,636 full-length LRR-malectin RLKs sequences were aligned using the muscle 5.1 software (*74*). Not trimmed alignment file was used in phylogenetic inference with FastTree 2.1.11 software (*75*). Based on the taxonomic tree, *S. muscicola* was the most basal species in this analysis. Therefore, the SM000010S04172, SM000162S02393 and SM000239S08065 proteins of *S. muscicola* were used to root the phylogenetic tree. Tree was visualized and annotated by the online website https://www.chiplot.online/.

### Statistical analysis of IGPs

Correlation analyses among LRR-malectin RLKs, IGPs, and primary gene models were performed using R 4.0.3, with the Pearson’s correlation test. The statistical significance of the difference between special lifestyle, habitats, or trophic types (SLHT) and non-SLHT species was assessed using unpaired two-sample t-test. Comparison of IGPs number between SLHT and non-SLHT species within the same order was performed using Wilcoxon rank-sum test. Statistical analysis results were visualized using the R package ggplot2.

### Preparations of polysaccharide solutions

Cellooligomers (CEL2, CEL3, CEL4), 3^1^-β-_D_-cellobiosyl-glucose (MLG43), XXXG (a hepta-saccharide hydrolysate of xyloglucan), mannohexaose (MAN6), xylotetraose (XYL4) were purchased from Megazyme. All the polysaccharides were dissolved in water or the buffer containing 10 mM Bis-Tris pH 6.0, 100 mM NaCl to prepare the 10 mM stock solution.

### Protein expression and purification

The ectodomains of IGP1 (residues 1-627), IGP4 (residues 1-635), IGPL1 (AT1G56120, residues 1-635) or their mutants with an engineered C-terminal 6xHis tag were cloned into a pFastBac1 vector using High-Fidelity PCR Mix (I-5™, TsingKe Biotech Co., Ltd, China). All proteins were expressed using the Bac-to-Bac baculovirus expression system (Invitrogen) with one litre of High Five cells (2.0×10^6^ cells/mL) infected with 30 mL recombinant baculovirus at 22 °C. The proteins were purified by Ni-NTA (Novagen) affinity chromatography from the supernatant harvested by centrifugation after 60 h post-infection. All proteins were further purified by size-exclusion chromatography (Hiload 16/600 Superdex 200 prep grade, GE Healthcare) in a buffer containing 10 mM Bis-Tris pH 6.0, 100 mM NaCl. The BAK1^LRR^ was expressed and purified as reported previously (*76*).

### Crystallization, data collection, structure determination and refinement

For crystallization, the purified IGP1^ECD^ and IGP4^ECD^ were concentrated to about 10 mg/mL. Crystals were generated by hanging-drop vapor-diffusion methods by mixing 1 μL of protein with 1 μL of reservoir solution at 18 °C. Diffraction quality crystals of apo-IGP1^ECD^ or apo-IGP4^ECD^ were obtained in the buffer containing 0.2 M potassium phosphate monobasic, 20% w/v polyethylene glycol 3,350 or 0.2 M Magnesium chloride hexahydrate, 0.1 M HEPES pH 7.5 25% w/v polyethylene glycol 3,350, respectively. IGP1^ECD^ was co-crystalized with CEL3 or CEL4 with a molar ratio of 1:10. Crystals of IGP1^ECD^-CEL3/4 complexes were both got in the 0.1M Sodium citrate tribasic dihydrate pH 5.0, 18% w/v polyethylene glycol 20,000 buffer. All crystal samples were flash frozen using the reservoir buffer plus 17% glycerol as the cryoprotectant. Diffraction datasets were collected at Shanghai Synchrotron Radiation Facility (SSRF) on beam line BL02U1, BL10U2, BL18U1 and BL19U1. Data were processed using the HKL2000 software package (*77*). The crystal structure of apo-IGP1^ECD^ was solved before Alphafold came out and thus determined by molecular replacement (MR) with PHASER (*78*) using the structure of ERECTA-like 1 (ERL1)^LRR^ (PDB code: 5xjo) and MALECTIN from *Xenopus laevis* (PDB code: 2k46) as searching models. The apo-IGP4^ECD^ structure was determined by molecular replacement (MR) with PHASER using apo-IGP1 as the template model. The model from MR was built with the program COOT (*79*) and subsequently subjected to refinement by the program Phenix (*80*) with excellent stereochemistry (**Table S5**). Structural figures were prepared using the programs PyMOL and COOT.

### Isothermal titration calorimetry assays

Binding affinities were measured using ITC200 (Microcal LLC) at 25 °C in a buffer containing 10 mM Bis-Tris, pH6.0 and 100 mM NaCl. The carbohydrate ligands (approximately 0.25 mM) were injected (20 × 2.0 μL) at 100 s intervals into the stirred (700 rpm) calorimeter cell (volume 250 μL) containing 0.025 mM IGP^ECD^ proteins. Measurements of the binding affinity of all the titration data were analyzed using the ORIGIN 7 software (MicroCal Software).

### LC–MS/MS analysis of IGP1 C591-C592 oxidation

To determine the oxidation status of C591 and C592, the IGP1 protein was expressed in Arabidopsis protoplasts (∼10 mL at a concentration of 2 × 10^5^/mL) for 16 h and treated with 10 uM CEL4 for 30 min. Protoplasts were then lysed with lysis buffer (50 mM HEPES-KOH, pH 7.5, 1 mM EDTA, 1% Triton X-100, 1% SDS and 1× complete protease inhibitors) and immunoprecipitated with GFP-trap beads (Sigma-Aldrich, USA). The immunoprecipitants were separated by 10% SDS-PAGE and stained with GelCode Blue Stain Reagent (Thermo Fisher, USA). The IGP1 bands were sliced, trypsin-digested, and oxidized-peptides were enriched for LC-MS/MS analysis using an Orbitrap QE LC-MS/MS system (Thermo Scientific, USA). The identified oxidized-peptides were manually inspected to ensure confidence in the oxidation site assignment.

### Plant materials and growth conditions

Arabidopsis plants used in this study are in the Col-0 background and were grown in soil at 22°C under 12 h/12 h light/dark photoperiod with 55% relative humidity for about 4 weeks. Arabidopsis T-DNA insertion lines *igp1-2* (SALK_016280C), *igp2/3-2* (SALK_144609C), *igp4-2* (SALK_005808C), *igp1l-2* (SALK_203752C) were obtained from the Arabidopsis Biological Resource Center (ABRC). *pIGP1::IGP1-FLAG/igp1-2*, *pIGP1::IGP1^W146A^-FLAG/igp1-2*, *pIGP1::IGP1^Y196A^-FLAG/igp1-2*, *pIGP1::IGP1^W220A^-FLAG/igp1-2*, *pIGP1::IGP1^W146A-Y196A-W220A (3M)^-FLAG/igp1-2*, *pIGP1::IGP1^C591A-C592A^-FLAG/igp1-2*, *pIGP1::IGP1^C591D-C592D^-FLAG/igp1-2*, and *pIGP1::IGP1^C591R-C592R^-FLAG/igp1-2* transgenic plants were generated using *Agrobacterium tumefaciens*-mediated floral dipping. Transgenic plants were screened by hygromycin (50 μg/mL) for pCAMBIA1300 vector, and confirmed by immunoblotting for protein expression.

The soybean Williams, which is susceptible to *Phy. sojae* P6497, was grown in plastic pots containing vermiculite at 25°C for 4 days in the dark. Etiolated seedlings were inoculated by pipetting 10 μL of the zoospore suspension on the hypocotyls, and maintained in a climate-controlled room at 25°C and 80% relative humidity in the dark.

*N. benthamiana* plants are routinely maintained in climate chambers at 19°C-22 °C for 4-6 weeks with a 14 h light/10 h dark photoperiod and LED lamps with a light intensity of ∼120-150 µmol m^−2^ s^−1^.

### Cellooligomers-induced immune assays

To measure oxidation burst, leaf discs (diameter 0.5 cm) were excised from 4-week-old Arabidopsis Col-0 or mutant seedlings and floated overnight in 200 μL of sterile distilled water in 96-well plates. The water was replaced with 200 μL of reaction buffer containing luminol/peroxidase (35.4 mg/mL luminol, 10 mg/mL peroxidase) and various cellooligomer treatments. Luminescence was quantified using a GLOMAX96 microplate luminometer (Promega, Madison, WI, USA). The experiments for MAPK activation were carried out as described previously (*81*). MAPK activation was determined by immunoblots with anti-phospho-p44/42 MAPK antibody (Cell Signaling #4370).

### Pathogens infection assays

For bacterial growth assays, *Pseudomonas syringae* pv. *tomato* DC3000 (*Pst* DC3000) strains adjusted to an optical density at 600 nm (OD_600_) of 0.0001 were infiltrated into the leaves of 4-week-old Arabidopsis Col-0 or mutant plants using a needleless syringe. For bacterial quantification, three leaf discs (6 mm diameter) per plant were harvested with a tissue punch 3 days post-inoculation (dpi). Samples were ground in 10 mM MgCl_2_, serially diluted (10, 100, 1000 and 10000 times), and 50 μL of each dilution was spread onto the selective NYG medium. The number of colonies (CFU) was calculated, and bacterial growth was represented as CFU cm^−2^ of leaf tissue.

*Phy. sojae* P6497 and *F. graminearum* were cultured in 10% V8 agar medium and PDA medium at 25°C in the dark for infection assays, respectively. *Phy. sojae* zoospores were diluted to a concentration of 100 zoospores/10 μL. Etiolated hypocotyls of soybean Williams were sprayed with different oligosaccharide solution. One day after spraying, *Phy. sojae* P6497 zoospore suspensions (100 zoospores) or *F. graminearum* mycelial plug (5 mm in diameter) were inoculated on the etiolated hypocotyls, and maintained at 25°C in the dark for 2 days, photographed and collected for biomass measurement. For oligosaccharides-induced resistance in potting assays, soybean plants at the 2-week growth stage were sprayed with different oligosaccharide solutions. One day after spraying, a *Phy. sojae* mycelial plug (5 mm in diameter) was inoculated onto the stem of each plant. The lodging rate of soybean plants was recorded 4 dpi.

For the soybean rust infection assay, urediniospores of *Pha. pachyrhizi* were suspended in a solution containing 0.1% Tween to prepare the spore suspension. Soybean plants at the 2-week growth stage were first sprayed with different oligosaccharide solutions 30 min before whole-plant spraying of the spore suspension. After seven days of incubation, disease incidence and biomass were recorded.

For the *Phy. capsici* infection assay in *N. benthamiana*, 4-week-old *N. benthamiana* leaves were sprayed with different oligosaccharide solution. One day after treatment, a 5-mm-diameter *Phy. capsici* mycelial plug was inoculated onto each leaf. Photos were taken and biomass was measured 2 dpi.

For the *Phy. capsici* infection assay in tomato, 4-week-old tomato leaves were collected and placed in trays. The leaves were first treated with an elicitor solution via spray application and incubated for 24 h to induce preliminary immune responses. Following this, a 4-mm diameter mycelial plug of *Phy. capsici* was inoculated onto each leaf. Photos were taken and biomass was measured 2 dpi.

For the inoculation assay with *B. graminis* (wheat powdery mildew), wheat seedlings at the coleoptile emergence stage were selected and transplanted into vermiculite-filled pots. The plants were cultivated in a controlled environment chamber maintained at 18°C and 60% relative humidity for 10-14 days to ensure uniform growth. Prior to pathogen challenge, the leaves of each pot were uniformly sprayed with 5 mL of an oligosaccharide solution and allowed to incubate for 24 h under the same conditions. Inoculation was subsequently performed by gently tapping leaves from heavily sporulating, diseased wheat plants above the treated seedlings, facilitating the even dispersal of *B. graminis* conidia onto the leaf surfaces. Following inoculation, the plants were returned to the controlled environment (18°C, 60% RH) for a 7-day period to allow for disease development before assessment.

### Genome-wide association study (GWAS) assays

A total of 349 wheat accessions were assembled to form an association panel for evaluating resistance to Fusarium crown rot (FCR) and leaf rust (LR). FCR severity was assessed 28 days after inoculation with the *Fusarium pseudograminearum* strain Fp-IV using a 0–9 disease index (DI) scale. For LR, adult plants were evaluated under field conditions in Baoding and Yuanyang following artificial inoculation. The maximum disease severity (MDS) of each accession was recorded when the susceptible control exhibited peak infection levels and used in subsequent analyses. All accessions were genotyped using the Wheat 660K SNP array by Beijing CapitalBio Technology Company (http://cn.capitalbio.com/), and GWAS was performed based on the obtained genotype data. Genotyping data quality control was performed using the PLINK software (version 1.9), with thresholds set to a minor allele frequency (-maf) of 0.02 and a missing genotype rate (-geno) of 0.1. GWAS was conducted in R using the GAPIT package, applying a mixed linear model that incorporated both population structure (PCA) and genetic relatedness (K matrix) (*82*). The kinship matrix was computed according to the VanRaden method. A modified Bonferroni correction approach (*83*) was applied to determine the significance threshold *P*, which was set at 10^−3^.

### Barley stripe mosaic virus (BSMV)-mediated gene silencing of *TaIGP1*

Virus-induced gene silencing (VIGS) was performed to silence *TaIGP1* (*TraesCS7A03G0083800*). A 178-bp fragment specific to the *TaIGP1* coding sequence was amplified from the wheat cultivar Zhuangguochun and cloned into the pSL038-1 vector to generate the BSMV_γ-*TaIGP1*_ construct. Plasmids containing the BSMV components (α, β, γ), the positive control BSMV_γ-PDS_ (targeting the phytoene desaturase gene), and the recombinant BSMV_γ-*TaIGP1*_ were linearized and used as templates for in vitro transcription. The resulting RNAs were mixed in a 1:1:1 ratio (α:β:γ or recombinant γ) and mechanically rub-inoculated onto the second leaves of seedlings of the BSMV-sensitive wheat cultivar Zhengmai9023 (*84*). Inoculated plants were kept in darkness for 24 h and then transferred to a growth chamber maintained at 23–25 °C with 60–80% relative humidity. At seven days post-virus inoculation (dpi), plants were challenged with the powdery mildew pathogen *B. graminis* isolate E09 by dusting with heavily sporulated colonies from infected donor plants. Powdery mildew resistance was assessed at two weeks post-inoculation, and leaf tissues were collected for subsequent physiological measurements. Total RNA was extracted from leaves for qRT-PCR analysis to determine the silencing efficiency of *TaIGP1*.

### Soybean hairy root transformation and *Phy. sojae* infection assays

For co-silencing *GmIGP1*, *GmIGP1L* and *GmIGP2* in soybean hairy roots, 200-bp DNA fragments of the indicated genes (*GmIGP1*, *GmIGP1L*, and *GmIGP2*) without predicted off-targets were designed via the Solanaceae Genomics Network (https://sol genomics.net) and were amplified from soybean Hefeng47 cDNA. The fragments were individually inserted into the pFGC5941-GFP vector, cloned in-frame with a Chalcone synthase linker from *Petunia hybrida*. Soybean seeds were cultivated at 25 °C the greenhouse for 6 days and the cotyledons were collected. Soybean cotyledons were surface sterilized with 10% (v/v) HClO for 20 min, then soaked in 75% (v/v) ethanol for 1 min, and washed with autoclaved dH_2_O five times. A small wound was made by cutting the lower epidermis of each sterile cotyledon, which was infected with a 10 mL suspension of *Agrobacterium rhizogenes* cells (K599) (OD_600_ = 0.5) carrying the indicated constructs. The soybean cotyledons were grown on Murashige and Skoog (MS) medium in 25 °C incubators without light. After 3 weeks, soybean transgenic hairy roots were screened by fluorescence stereomicroscopy (Nikon, SMZ25) using the GFP filter and further confirmed by qRT-PCR analysis. The hairy roots were inoculated with *Phy. sojae* mycelium, and the oospores and biomass were analyzed at 48 hpi.

### Protoplast-transient assay

3-week-old Arabidopsis leaves were used for protoplast isolation. Subsequently, 100 μg of the IGP1-GFP-pM999 plasmid was used to transform 1 mL of protoplasts, followed by incubation for 16 h. The total proteins were isolated with 100 μL of extraction buffer (50 mM Tris-HCl pH 7.5, 150 mM NaCl, 10 mM EDTA pH 8.0, 0.1% Triton X-100, 0.2% IGEPAL CA-630, 10 mM dithiothreitol, 1 mM phenylmethylsulfonyl fluoride, 1 × protease inhibitor cocktail). The samples were vortexed vigorously for 30 s and then centrifuged at 15,000 × *g* for 15 min at 4 °C. The supernatant was incubated with α-GFP agarose beads for 2 h at 4°C with gentle shaking. The conjugated beads were collected by centrifugation at 500 × g at 4°C for 3 min and washed three times with 500 μL washing buffer (50 mM Tris-HCl pH 7.5, 150 mM NaCl). Bound proteins were released from the beads by boiling in SDS/PAGE sample loading buffer and analyzed by Western blotting (WB) with an α-GFP antibody.

### Agroinfiltration of *N. benthamiana*

*Agrobacterium tumefaciens* strains harboring distinct expression constructs were cultured overnight at 28°C in LB medium. The bacterial cells were subsequently pelleted by centrifugation at 3,600 g for 5 min, followed by three washes with ATTA infiltration buffer (10 mM MgCl_2_, 10 mM MES, pH 5.7, 100 nM acetosyringone). After resuspension in the same buffer, the cultures were conditioned by incubation for 2 h at room temperature to induce virulence gene expression. Prior to infiltration, the optical density at 600 nm (OD_600_) of each bacterial suspension was adjusted to standardize the inoculum: OD_600_ = 1.0 for strain carrying the pBIN-GFP-IGPs^ECD^ vector, and OD_600_ = 0.3 for the strain carrying the P19 silencing suppressor. The adjusted suspensions were then co-infiltrated into the abaxial side of leaves from 5- to 6-week-old *N. benthamiana* plants using a needleless syringe.

### MAL-PEG2000 assays

To assess the direct oxidation of recombinant protein thiol groups by hydrogen peroxide (H_2_O_2_), reactions were carried out in a 100 µL volume containing 2.5 µM of IGP1 protein in Buffer I (25 mM Tris-HCl, pH 7.5, 100 mM NaCl, 10 mM MgCl_2_, and 1 × complete protease inhibitors) supplemented with specified concentrations of H_2_O_2_. The mixture was incubated at room temperature for defined time periods (*37*). Following oxidation, residual H_2_O_2_ was removed using Zeba Micro Spin desalting columns (Thermo Fisher, #89883) to terminate the reaction. The eluted protein was subsequently incubated with 20 mM methoxy PEG maleimide (MAL-PEG2000; JenKem, #A3124-1) for 1 h at room temperature in the dark, allowing the maleimide group to covalently block any remaining free thiol groups. The modified proteins were then analyzed by SDS-PAGE. Since each conjugated MAL-PEG2000 moiety (average molecular weight ∼2,000 Da) introduces a measurable increase in apparent molecular mass, a discernible band shift between oxidized (unmodified) and reduced (PEGylated) protein states can be visualized, enabling the differentiation of their redox status based on electrophoretic mobility

### Quantitative RT-PCR analysis

Total RNA was extracted from 4-week-old Arabidopsis Col-0 seedlings or 2-week-old soybean Williams seedlings using RNA-easy TM Isolation Reagent (Vazyme, Cat. No. R701-01) pre-infiltrated with controls (dH2O) or different oligosaccharide solution. 2 μg total RNA was subjected to synthesis of cDNA using All-in-One 5×RT MasterMix (abm, Cat. No. G592), and quantitative (q) RT-PCR was performed with BlasTaq 2×qPCR MasterMix (abm, Cat. No. G891) on the BIO-RAD CFX96TM real-time system. For data analyses, the 2^−..6..6^Ct method was used to calculate the relative gene expression and the mRNA level of *ACT2* (AT3G18780) or *GmCYP2* (Glyma.12G024700) was used as an internal reference.

---

**fig. S1.**
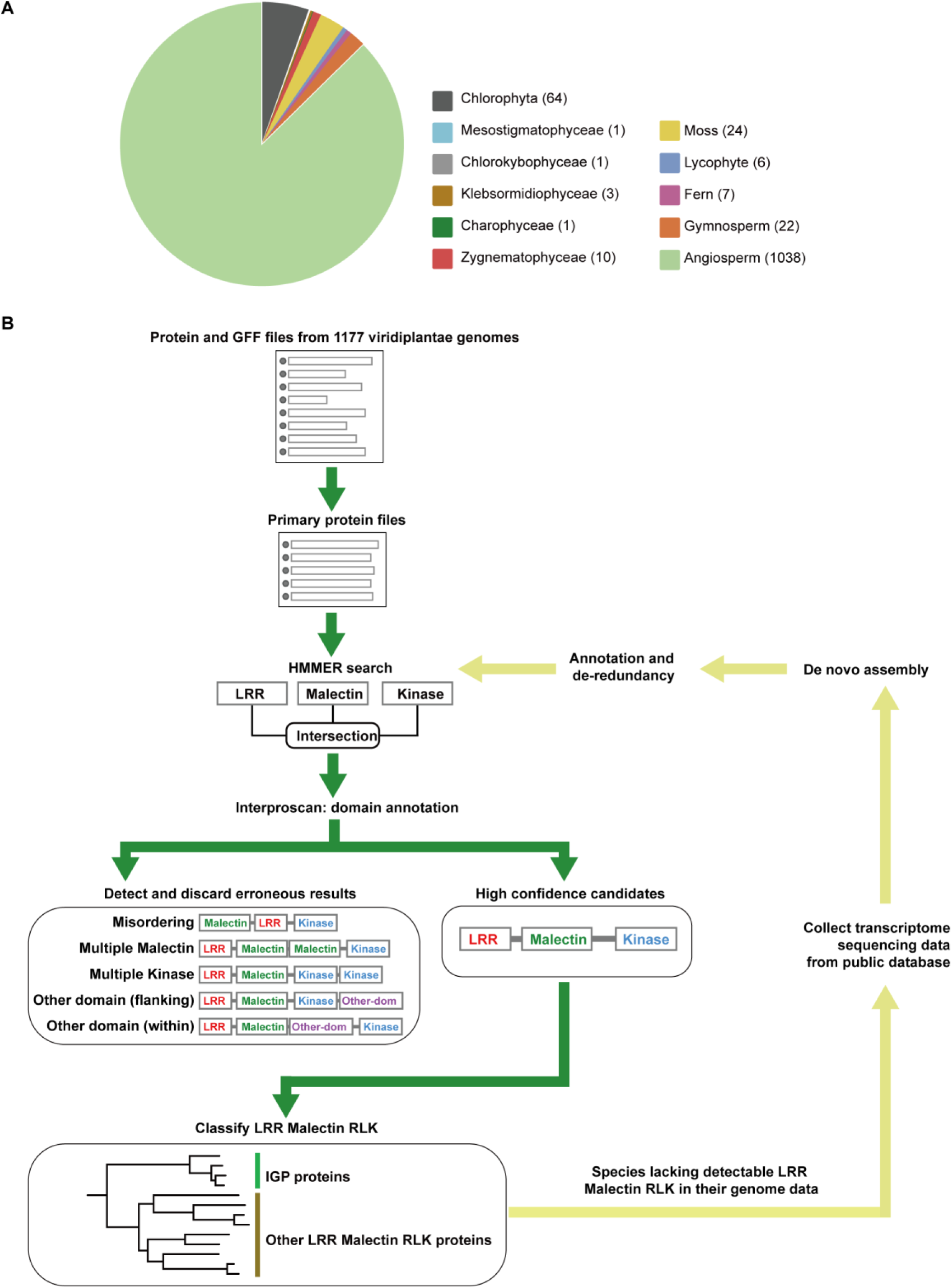
Identification pipeline for LRR-Malectin RLKs. (**A**) Summary statistics of the species genomes included in this study. (**B**) Schematic workflow for identifying LRR-Malectin RLKs.

**fig. S2.**
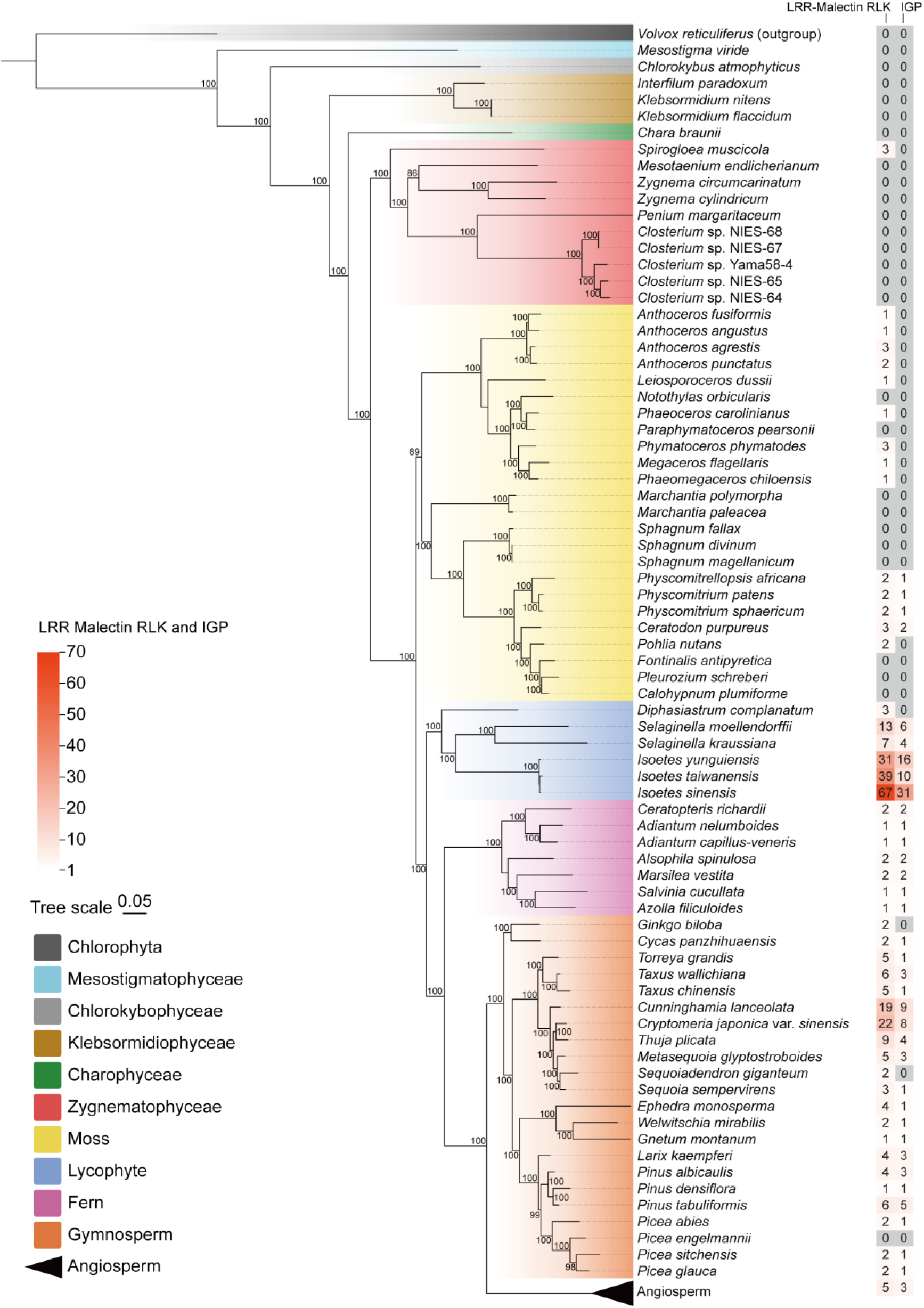
Analysis of LRR-Malectin RLKs in species tree. Phylogenetic tree of 77 species. The tree was constructed based on 67 single-copy orthologs. Algae, mosses, ferns, lycophytes, gymnosperms and angiosperms (*Amborella trichopoda*) were marked with different ground colors. The green algae, *Volvox reticuliferus*, was selected as outgroup. The number of LRR-malectin RLKs and IGPs are annotated to the right.

**fig. S3.**
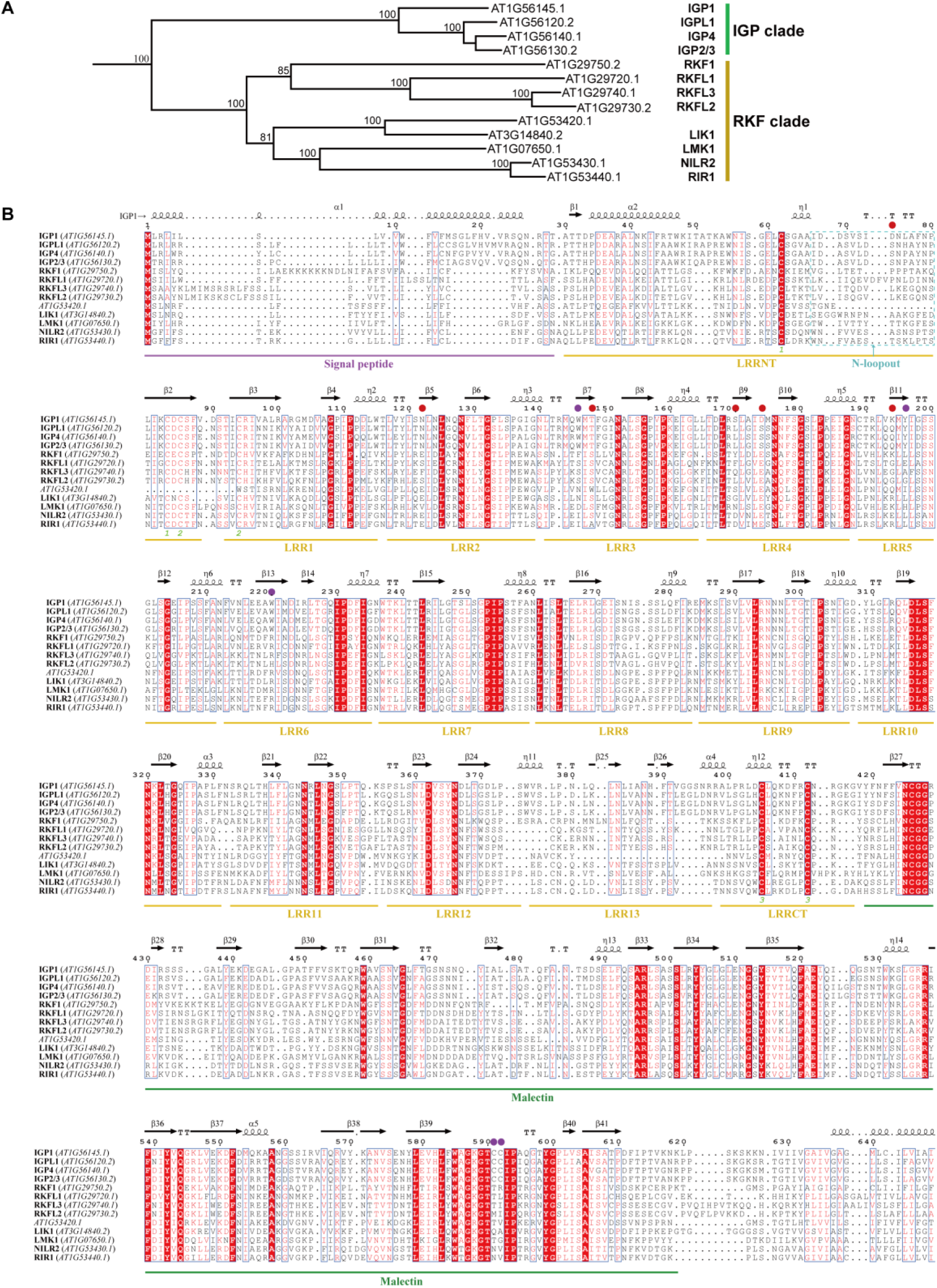
Phylogenetic tree and structure-based sequence alignment of Arabidopsis LRR-Malectin RLKs. (**A**) Phylogenetic tree of Arabidopsis LRR-Malectin RLKs. LIK1: LysM RLK1-interacting kinase 1. LMK1: LRR with Malectin RLK1. RIR1: Remorin-interacting receptor 1. NLIR2: Nematode-Induced LRR-RLK 2. RKF1: RLK in Flowers 1. RKFL1-3: RKF-like 1-3. (**B**) Structure-based sequence alignment of Arabidopsis LRR-Malectin RLKs. Regions of LRRNT/CT, N-loopout, LRR1-13 and malectin are underlined as indicated. Important residues for IGP1 interaction with CELs are marked with purple (for CC-WYW motif) and red dots. No potential polysaccharide-binding pocket was detected in RKF clade.

**fig. S4.**
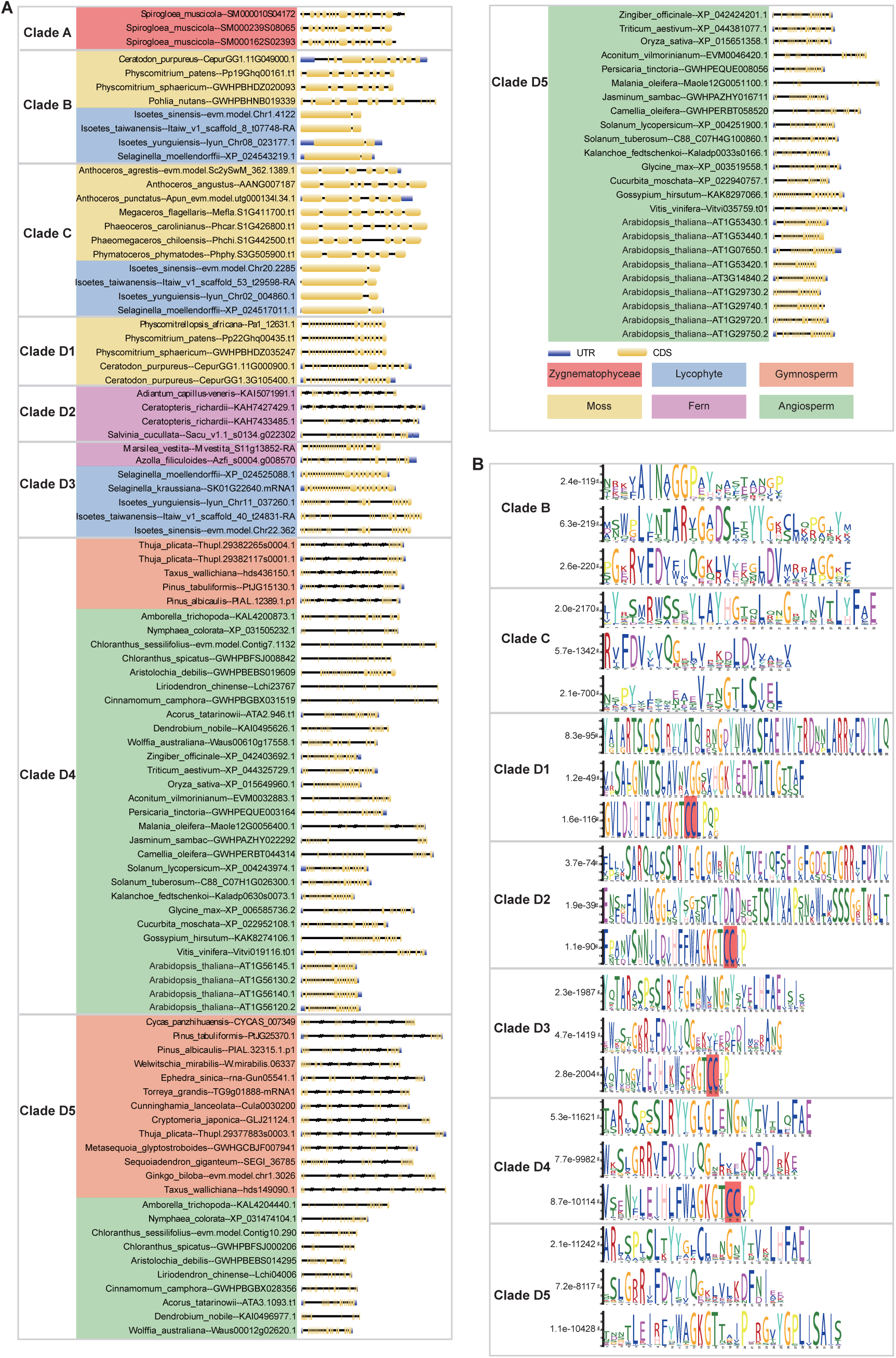
Gene structure and conserved motifs analysis of LRR-Malectin RLKs. (**A**) Gene structure of 108 structurally complete *igps*. Different plant groups are highlighted with distinct background colors, Zygnematophyceae-red, mosses-yellow, lycophytes-blue, ferns-purple, gymnosperm-orange, angiosperm-green. Gene structure information was extracted from the GFF files. (**B**) Conserved motifs of malectin domain in clades B-D. The sequence logos of conserved motifs are displayed. E-value of each motif is annotated.

**fig. S5.**
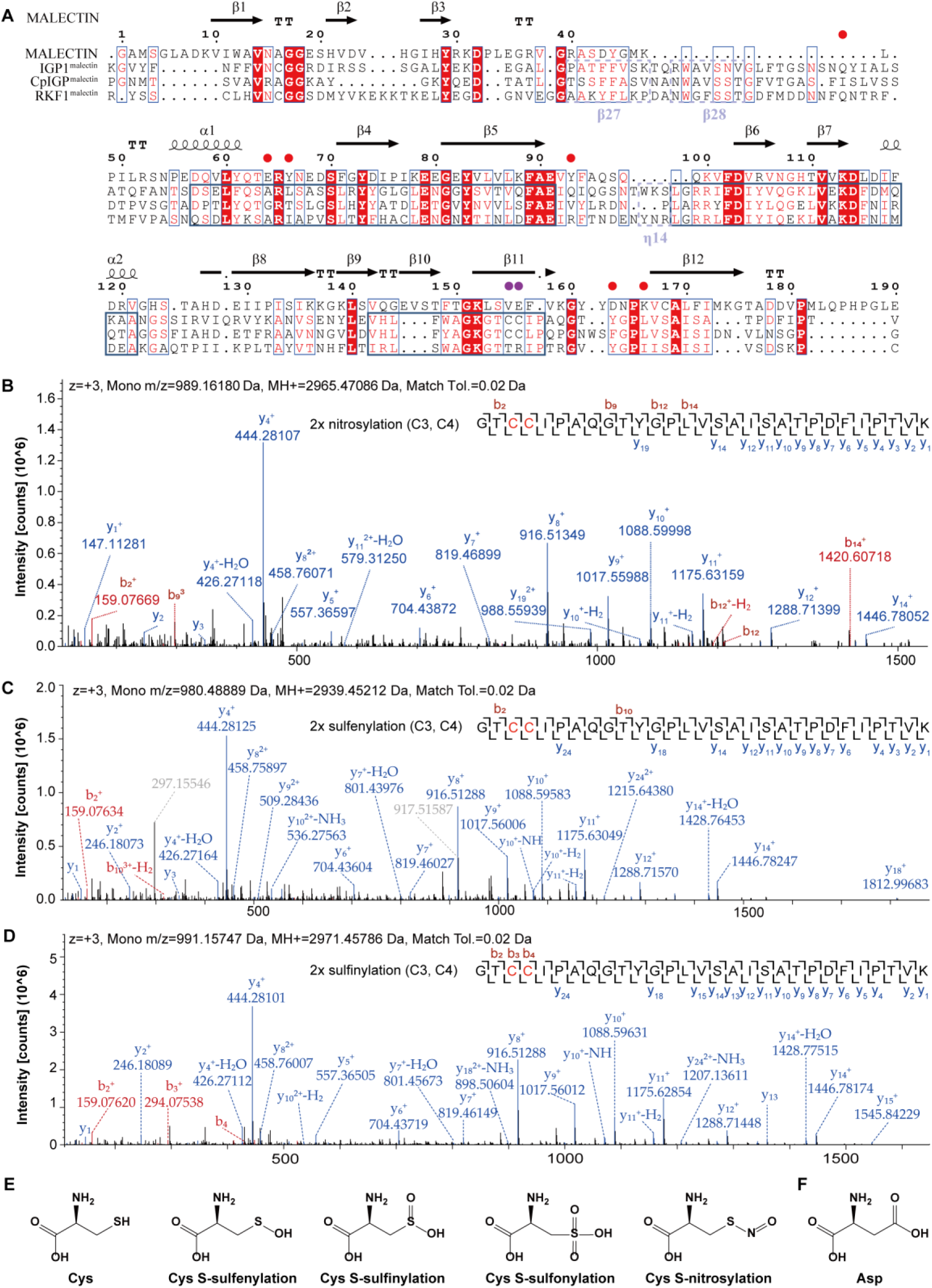
The CC motif in IGP1^Malectin^ can be oxidic/nitrosyl-modified. (**A**) Structure-based sequence alignment of MALECTIN, CpIGP1^malectin^ (subclade D1), IGP1^malectin^ (subclade D4), and RKF1^malectin^ (subclade D5). Cp, *Ceratodon purpureus*. The identified conserved motifs of malectin domain in clades B-D (**fig. S3B**) are framed in dark blue. Purple dots mark the unique CC motif in IGP1^malectin^ and CpIGP^malectin^. Red dots mark the residues crucial for MALECTIN recognizing nigerose. (**B-D**) LC– MS/MS spectrum recorded on the [M+3H]^3+^ ionof the IGP1 CC peptide harboring oxidative modification. Predicted b- and y-type ions are listed. Detected ions are labeled in the spectrum and indicate that C3 and C4 of the peptide are nitrosyl-(**B**), sulfenyl-(**C**) and sulfinyl-(**D**) modified, respectively. (**E**) Structural formula of Cys, Cys with S-sulfenylation (-SOH), sulfinylation (-SO_2_H), sulfonylation (-SO_3_H) and nitrosylation (- SNO). (**F**) Structural formula of Asp.

**fig. S6.**
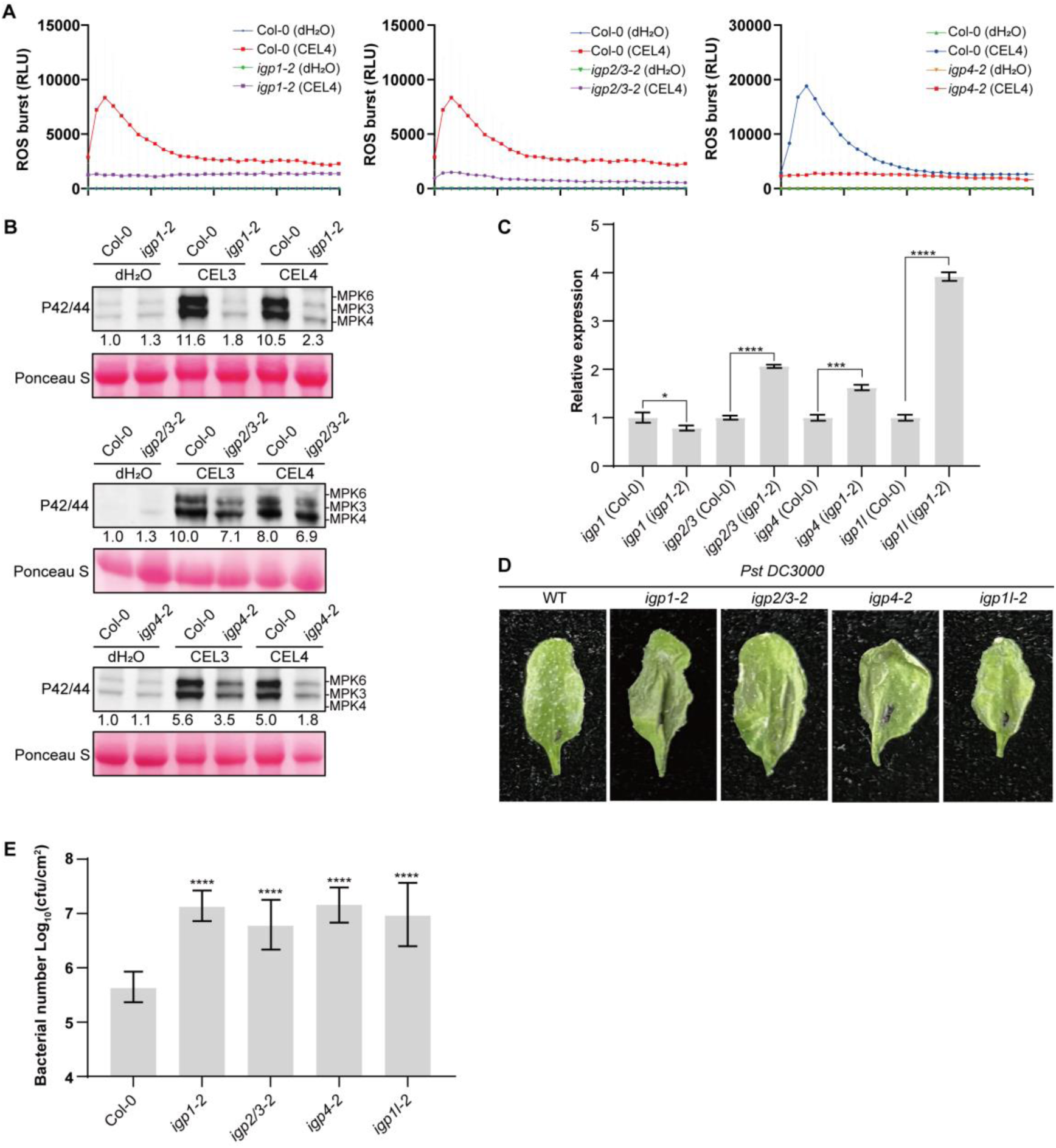
IGPs positively regulate cellooligomers-induced immune responses and plant disease resistance. (**A** and **B**) ROS burst (**A**) and MAPK activity (**B**) induced by cellooligomer in *igp1-2*, *igp2/3-2*, and *igp4-2* mutants. Phosphorylation of MAPK was detected using an anti-p42/p44 antibody. Ponceau S staining of the blot was used to verify equal protein loading. (**C**) Detection of *IGPs* expression levels in *igp1-2* mutant by qRT-PCR. (**D**) Visual symptoms 2 dpi with *Pst* DC3000 on the leaves of *igps* mutants. Representative photographs are shown. (**E**) Increased susceptibility to *Pst* DC3000 in *igps* mutants. Four-week-old soil-grown plants were hand-inoculated with *Pst* at 5×10^5^ cfu/mL. Data represent mean ± SEM (*n* = 6).

**fig. S7.**
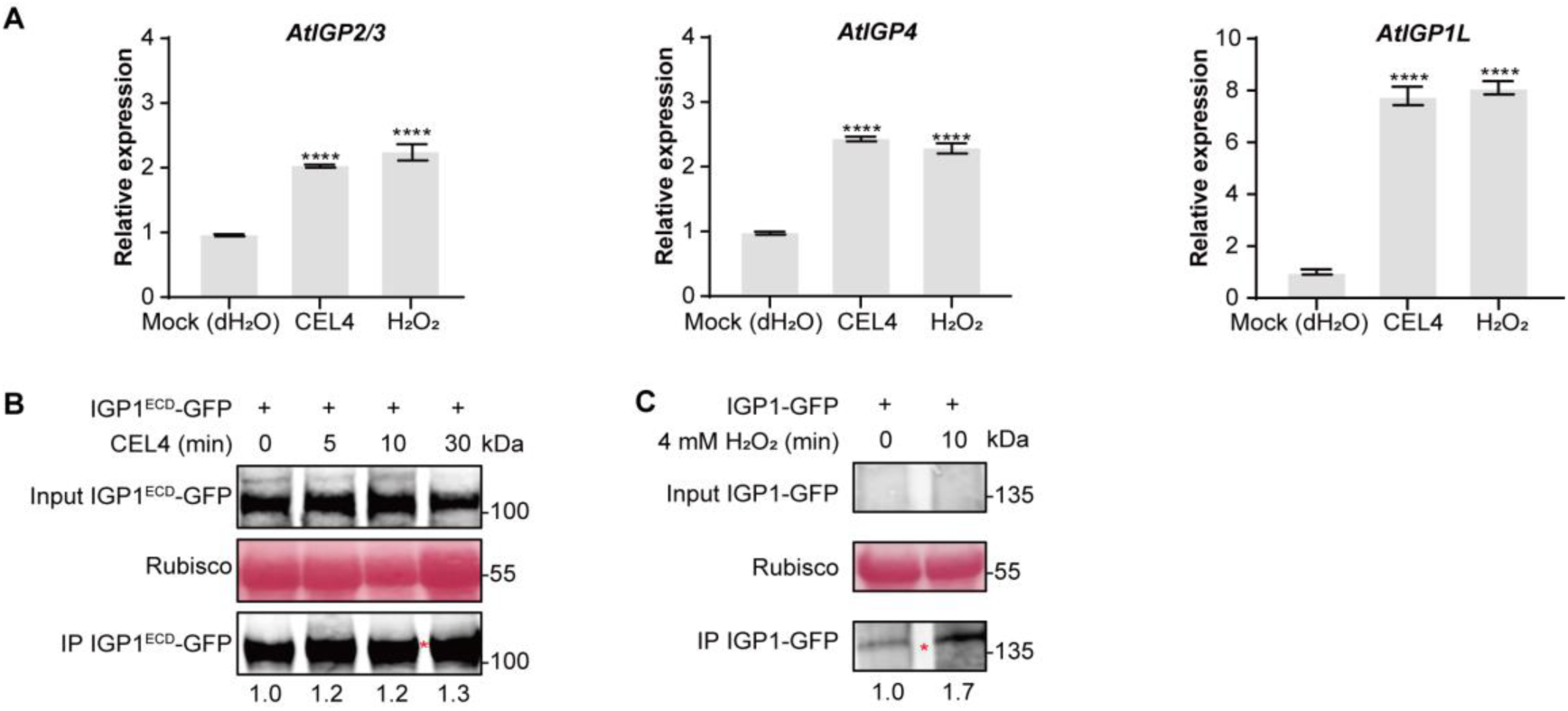
Cellooligomers and H_2_O_2_ induce the expression and protein accumulation of IGPs. (**A**) CEL4 and H_2_O_2_ induced *AtIGP*s expression. qRT-PCR was performed to quantify the expression level of *AtIGP2/3*, *AtIGP4*, and *AtIGP1L* in Arabidopsis after treatment with 10 µM CEL4 and 20 µM H_2_O_2_ for 40 minutes with dH_2_O as negative control. (**B**) Detection of IGP1^ECD^ protein expression in *N. benthamiana* following CEL4 treatment. The IGP1^ECD^-GFP protein was transiently expressed in *N. benthamiana* via *Agrobacterium*-mediated infiltration. Leaves expressing the protein were treated with 10 µM CEL4 for different durations (0, 5, 10, 30 min). The band intensity was quantified using ImageJ software. (**C**) Detection of IGP1 protein expression in *N. benthamiana* following H_2_O_2_ treatment. The transient expression of IGP1-GFP in *N. benthamiana* followed by treatment with 4 mM H_2_O_2_ for 10 minutes, and subsequent detection of IGP1-GFP protein expression levels.

**fig. S8.**
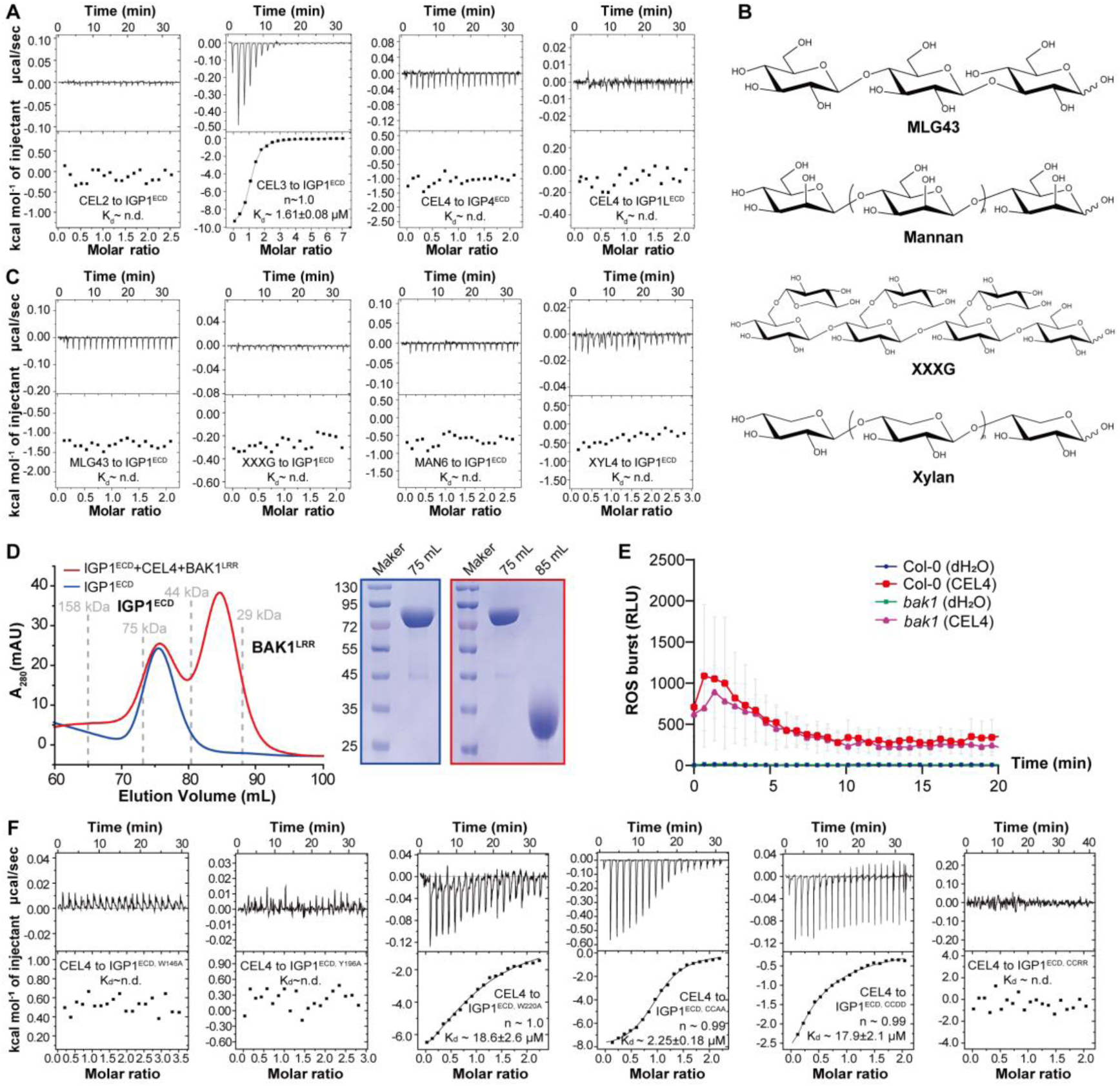
IGP1 specifically interacts with cellooligomers (DP≥3) and does not heterodimerized with BAK1. (**A**) Quantification of the binding affinity of CEL2/3-IGP1^ECD^, CEL4-IGP1^ECD^/IGPL1^ECD^ at pH 6.0 by ITC assays. The binding constants (Kd values ± fitting errors) and stoichiometries (n) are indicated. n.d.: no detectable binding. (**B**) Structural formula of hemi-celluloses tested in this study. MLG43: 3^1^-β-_D_-cellobiosyl-glucose. XXXG: a hydrolysate of xyloglucan. Xylan: polymer of β-1,4-_D_-xyloses; Manan: polymer of β-1,4-_D_-mannoses. (**C**) Quantification of the binding affinity of MLG43, XXXG, mannohexaose (MAN6), xylotetraose (XYL4) with IGP1^ECD^ at pH 6.0 by ITC assays. (**D**) Gel filtration profiles of IGP1^ECDD^ and a mixture of IGP1^ECD^, CEL4, BAK1^LRR^ at pH 6.0 100 mM NaCl shown in different colors. A280 (mAU), micro-ultraviolet absorbance at the wavelength of 280 nm. Coomassie blue staining of the gel filtration fractions the following SDS-PAGE shown on the right. CEL4 did not induce IGP1^ECD^ homodimerization nor IGP1^ECD^-BAK1^LRR^. (**E**) ROS burst induced by CEL4 in *bak1* mutant with dH_2_O as negative control. (**F**) Mutagenesis of IGP1-cellooligomer complex. Quantification of the binding affinity of CEL4-IGP1^ECD, W146A, Y196A, W220A, CCAA, CCDD or CCRR^ at pH 6.0 by ITC assays. In **A**, **C**-**F**, experiments were repeated three times with similar results.

**fig. S9.**
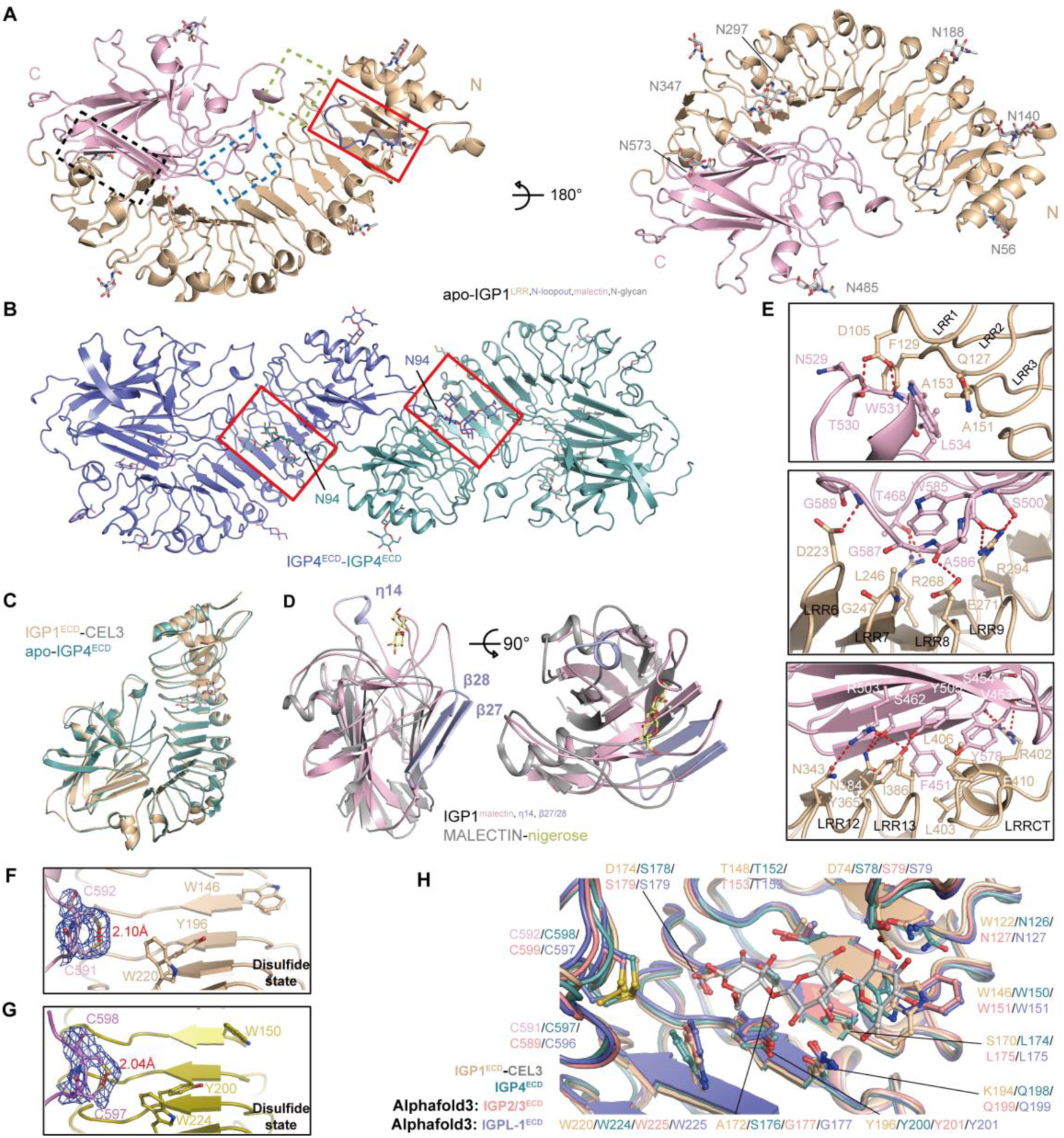
The LRR-malectin bipartite structure of IGPs. (**A**) Cartoon diagrams of apo-IGP1^ECD^ in two orientations. The LRR-malectin interface is framed in palegreen/blue/black dash lines. N and C represent N and C termini, respectively. Distributions of 7 N-glycans are colored in grey stick. (**B**) Two IGP4 N94-glycosylations mutually packs into the ligand-binding pocket of the other IGP4 molecules in one asymmetry unit of apo-IGP4^ECD^ structure. (**C**) Structural superposition of IGP1^ECD^-CEL3 and apo-IGP4^ECD^. r.m.s.d. of 0.710 Å compares the corresponding Cα atoms. (**D**) Structural superposition of IGP1^Malectin^ and MALECTIN-nigerose (PDB: 2k46). r.m.s.d. of 4.518 Å compares the corresponding Cα atoms. The extra structural elements β27/28 and η14 of IGP1^Malectin^ are depicted in lightblue. (**E**) Interaction details of LRRs and Malectin. Exhibition are important residues for interaction. Top, η14 in IGP1^Malectin^ interact with IGP1^LRR^. Middle, loops connecting β36, β37 in IGP1^Malectin^ interact with IGP1^LRR^. Bottom, β27, β28 in IGP1^Malectin^ interact with IGP1^LRR^. The color codes keep in same as indicated. (**F** and **G**) Electron density (2Fo-Fc) around C591-C592 of apo-IGP1^ECD^ (**F**)/C597-C598 of apo-IGP4^ECD^ (**G**) contoured at 1.5 sigma. The atom distances between the Cys-pair are labeled in red. (**H**) A conserved cellooligomer-perception pocket found in IGP1,2/3,4,L1. Structural comparison of ligand-binding pockets in IGP1,2/3,4,L1. Important residues for interaction are shown.

**fig. S10.**
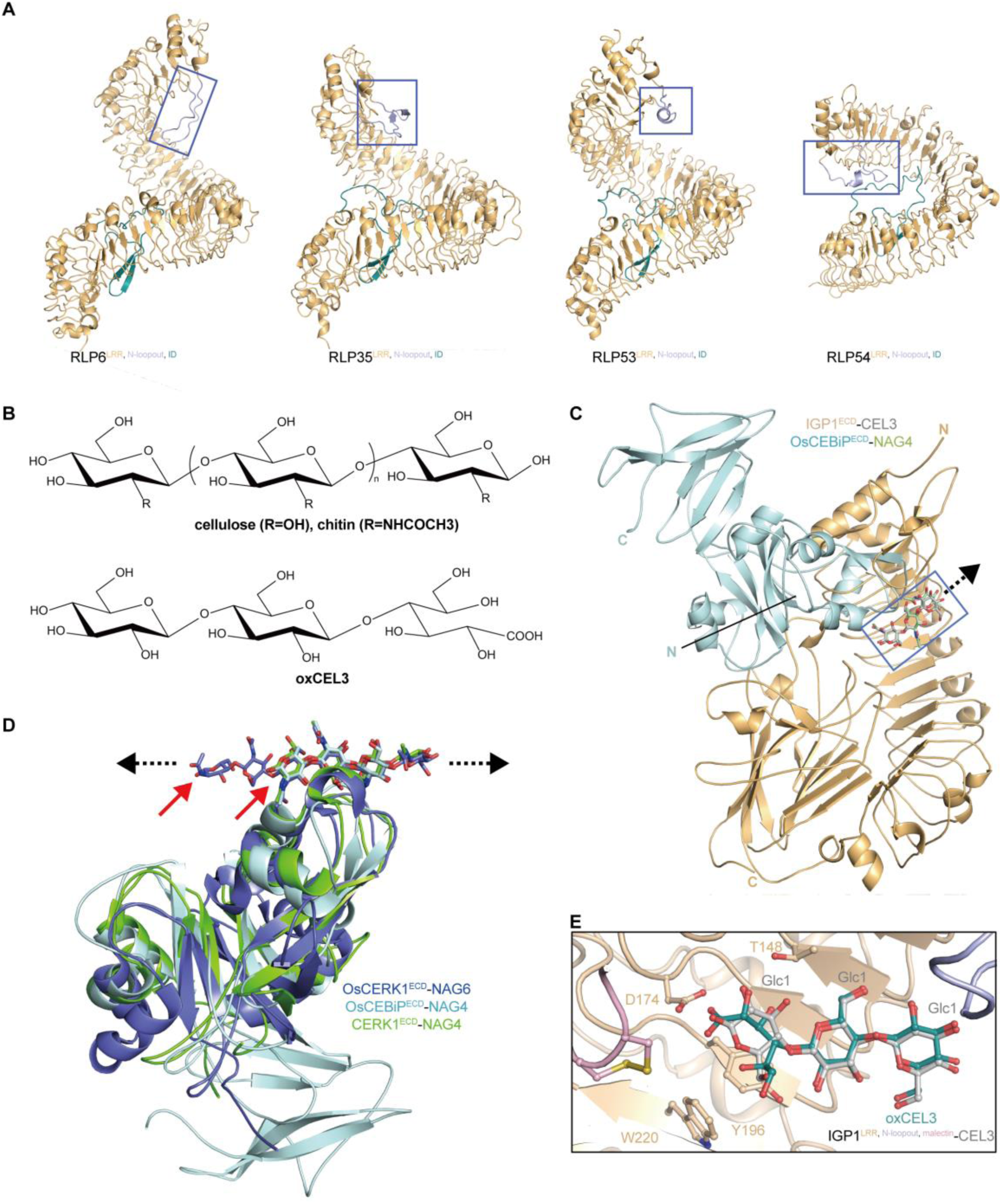
The N-loopout region of IGP1 and reducing end of cellooligomer is crucial for IGP1-cellooligomer interaction. (**A**) Alphafold 3-modeled structures of Arabidopsis RLP6/35/53/54. The N-loopout regions are framed in purple. (**B**) Structural formula of cellulose/chitin (upper) and OxCEL3 (bottom). Chitin (a homopolymer of N-acetyl glucosamine, NAG) is a derivative of cellulose. OxCEL3 is the oxidation products of CEL3 by CELLOXs (*49, 50*). (**C**) Structural superposition of IGP1^ECD^-CEL3 and OsCEBiP^ECD^-NAG4 (PDB: 5jce, only 3 NAG residues of NAG4 can be observed in the structure) based on the orientation of CEL3 and NAG4. N and C represent N and C termini, respectively. Blue frame highlights aligned oligosaccharides. Black arrows indicate the unidirectional extension of cellooligomer. (**D**) Structural superposition of OsCERK1^ECD^-NAG4 (PDB: 7vs7), CERK1^ECD^-NAG4 (PDB: 4ebz), OsCEBiP^ECD^-NAG4 based on the orientation of chitooligosaccharides. Red arrows indicate the reducing end of NAG4/6. Black arrows indicate the bidirectional extension of chitooligosaccharides. (**E**) Structural superposition of oxCEL3 with IGP1^ECD^-CEL3 complex shows steric incompatibilities.

**fig. S11.**
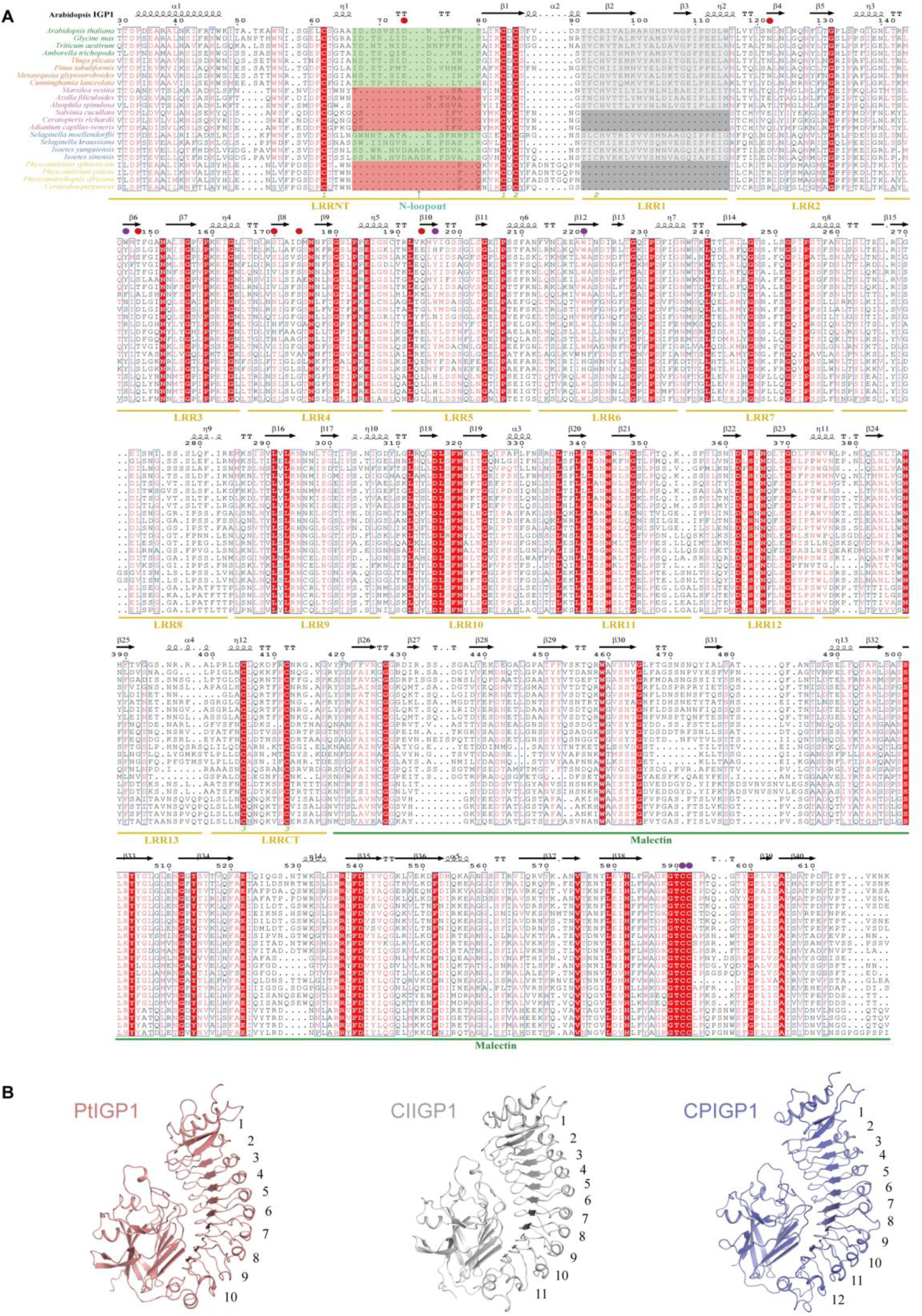
Structural conservation and divergence of plant IGPs. (**A**) Structure-based sequence alignment of IGP sequences from different plant groups. Species names from different plant group are labelled in different font colors: angiosperms-green, gymnosperms-orange, ferns-purple, lycophytes-blue, and mosses-yellow. N-loopout regions are highlighted with green background. For species lacking the N-loop, the corresponding sequence region is highlighted with a red background. All moss species and some fern species possess one fewer LRR compared to other plant. The corresponding regions are highlighted in dark grey. Important residues for IGP interaction with CELs are marked with purple (for “CC-WYW” motif) and red dots. (**B**) Representative structures of IGP^ECD^ with 10/11/12 LRRs modeled by Alphafold 3. LRRs are numbered in order from N- to C-terminus. PtIGP1, *Pinus tabuliformis* ptjg150001; ClIGP1, *Cunninghamia lanceolate* Cula0004975.1; CpIGP1, *Cunninghamia lanceolate* CepurGG1.3G105400.1.

**fig. S12.**
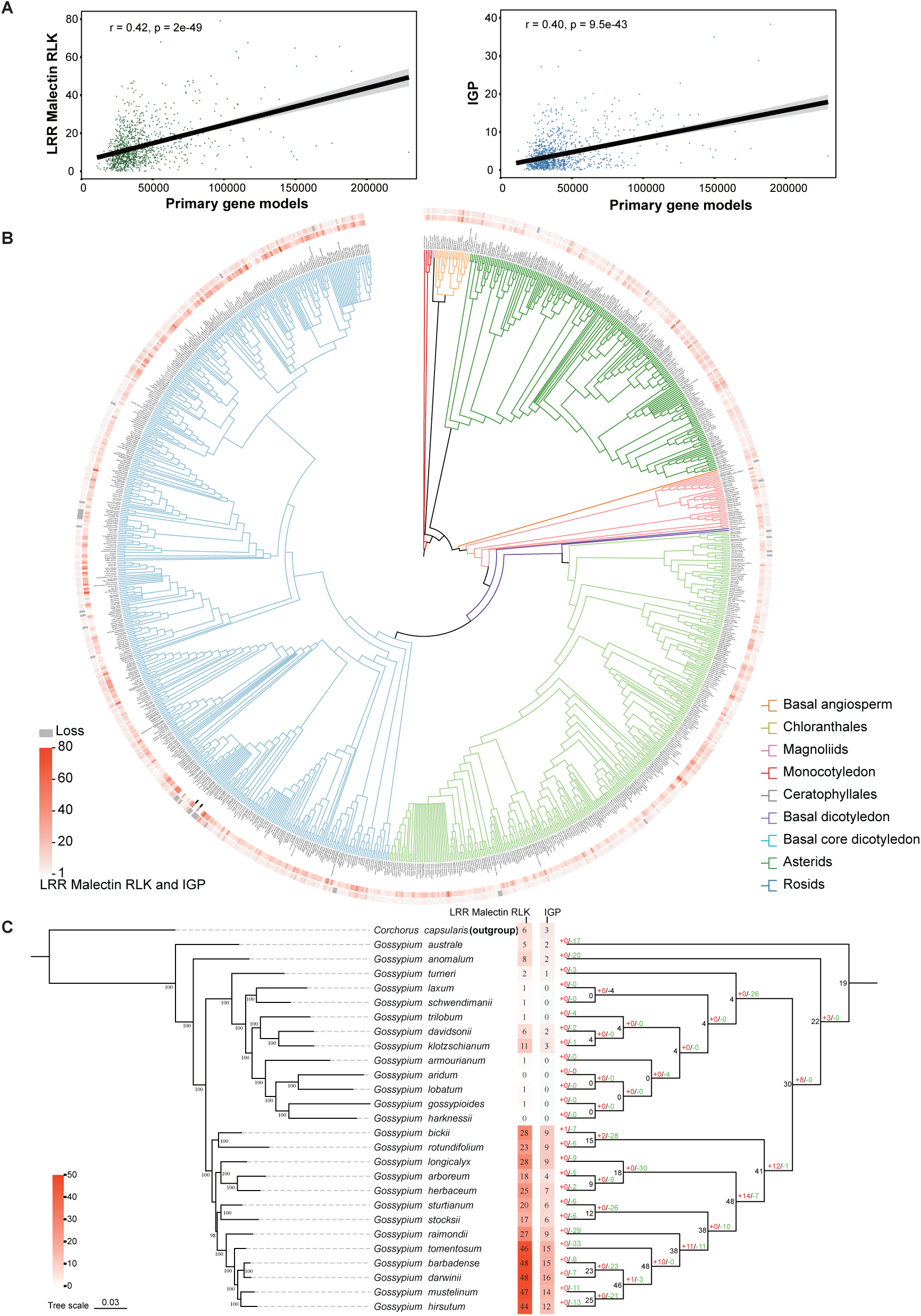
Evolution of angiosperm IGPs is shaped by genomic variation, lineage-specific expansions and contractions. (**A**) Correlation analysis reveals a moderate association between LRR-Malectin RLKs (left)/IGPs (right) and the total number of protein-coding genes. Pearson correlation coefficients and p-values are annotated. The black line represents the linear regression trend, and the grey shading indicates the confidence interval. (**B**) Evolutionary dynamics of LRR-Malectin RLKs and IGPs among 1,038 angiosperms. The inner ring represents the topology of the phylogenetic tree, with different clades distinguished by branch colors. Full species names are annotated at the corresponding leaf nodes. The two outer rings display heatmaps representing the number of LRR-Malectin RLKs and IGPs, respectively. Grey indicates the absence of LRR-Malectin RLKs or IGPs. (**C**) An example of lineage-specific expansion and contraction of IGPs from the genus Gossypium. Left: Phylogenetic tree of Gossypium species constructed from 2,390 single-copy orthologous genes, with *Corchorus capsularis* used as the outgroup. Right: Expansion and contraction analysis of IGPs, with expansions shown in red and contractions in green. The inferred number of IGPs at each ancestral node is indicated in black.

**fig. S13.**
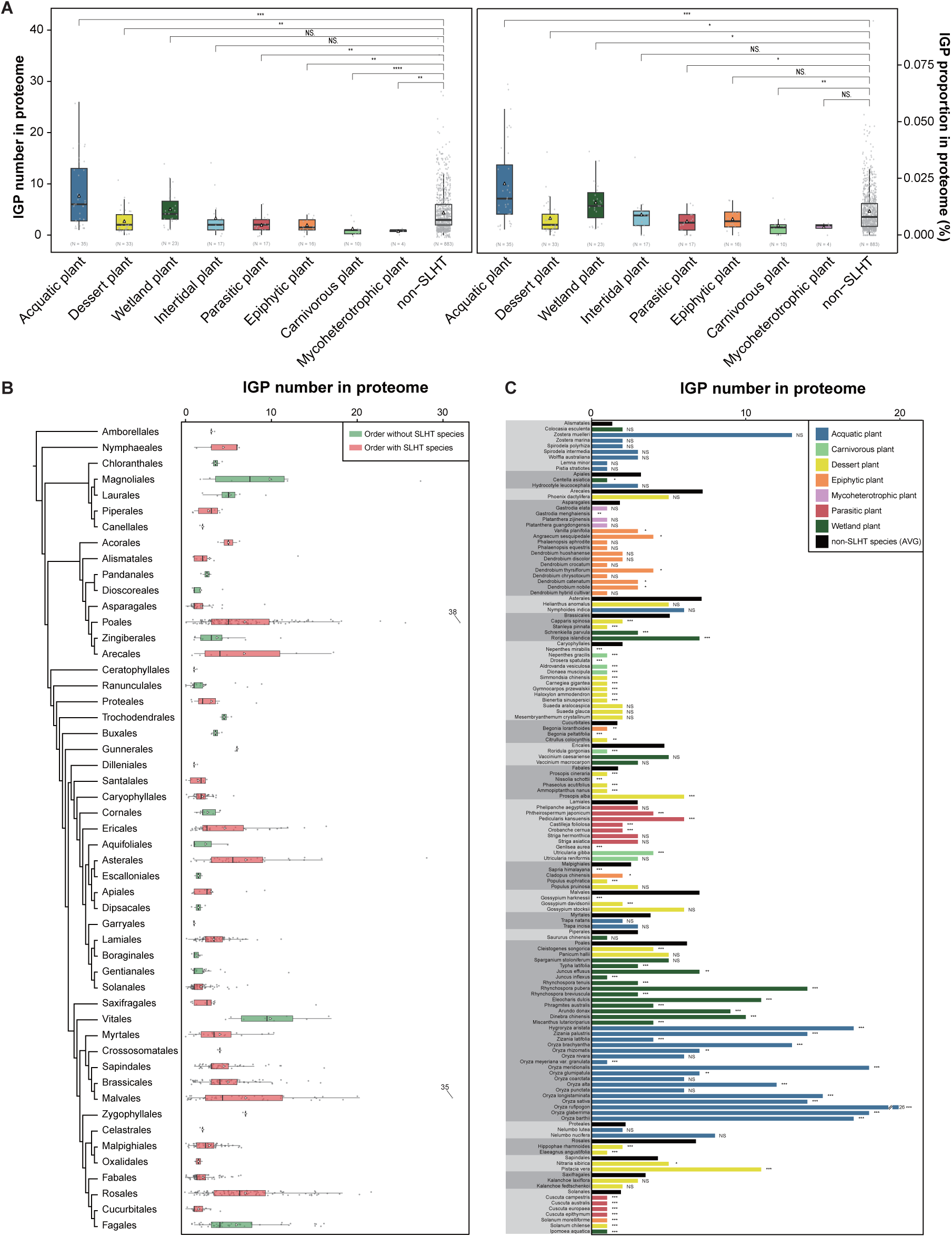
Evolution of angiosperm IGPs is shaped by ecological adaptation. (**A**) Comparative analysis of IGPs between species adapted to special lifestyle, habitats, or trophic types (SLHT) and non-SLHT species. The copy number of IGPs is increased in aquatic and wetland plants but reduced in desert species. Left: based on number of IGPs; right: based on percentage of IGPs among primary gene models. Significance levels are denoted as NS, *, **, and ***. (**B**) Distribution of the IGP numbers in each angiosperm order with or without SLHT species. The phylogeny at the order level on the left was based on APG IV. The number of IGPs in different orders is displayed using boxplots. Orders containing SLHT species are highlighted in red, while those without SLHT species are highlighted in green. No significant difference was found between orders that include SLHT species and those that did not. (**C**) Comparison of IGP numbers between SLHT and non-SLHT species within the same order. Each order is shaded alternately with light and dark grey backgrounds. Order names are presented in bold font. In the bar plot, black bars represent the average number of IGPs in non-SLHT species within each order, while other colors correspond to the IGP numbers in aquatic, desert, wetland, intertidal, parasitic, epiphytic, carnivorous, and mycoheterotrophic plants, respectively. Within each order, statistical significance of difference in IGP numbers between SLHT and non-SLHT was tested by Wilcoxtest method. In most orders, the number of IGPs in SLHT species significantly differs from the average number observed in non-SLHT species within the same order (p-value < 0.05). These findings suggest that both environmental selection and lineage-specific evolutionary events may have jointly contributed to the variation in IGP copy numbers.

**fig. S14.**
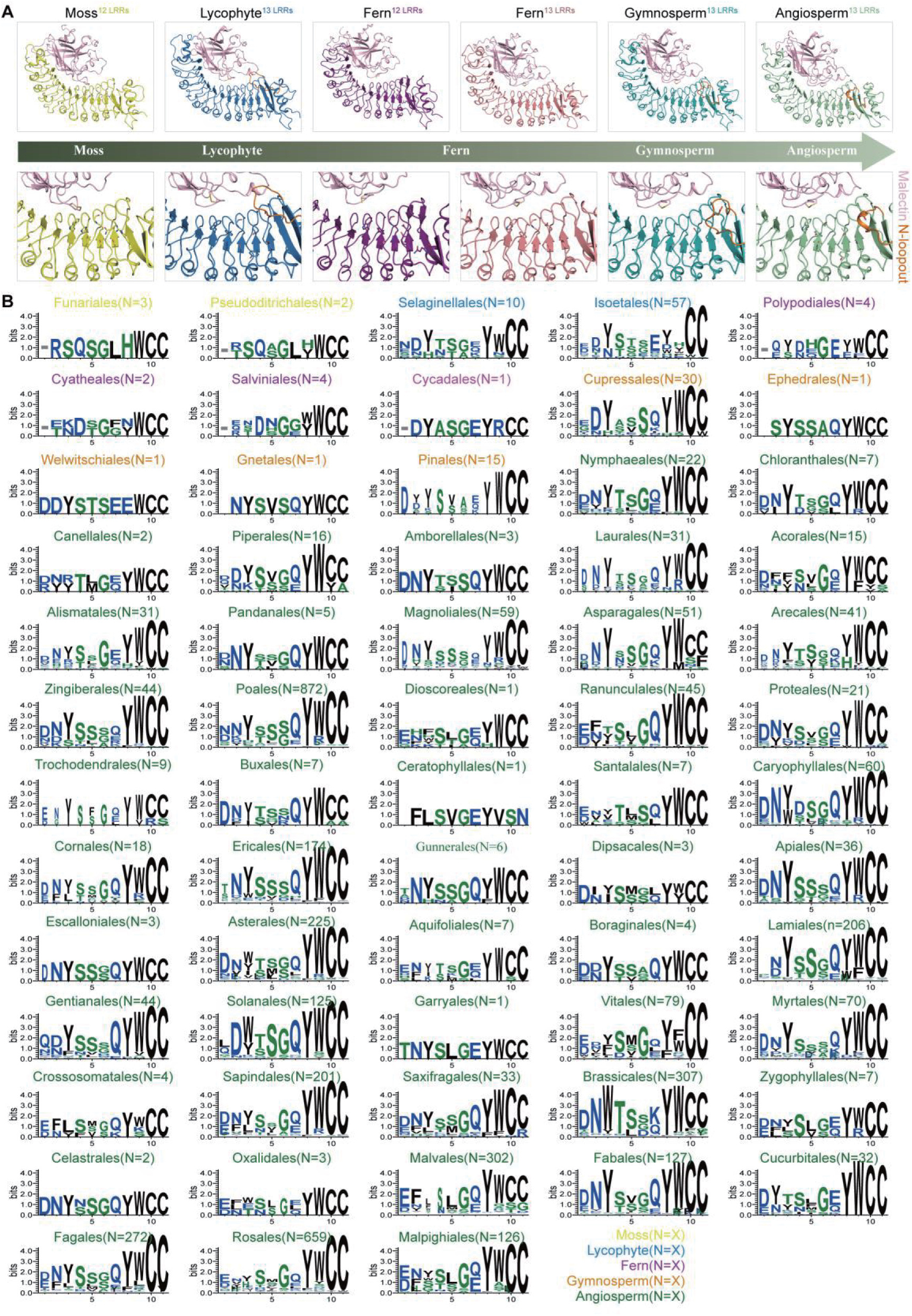
Evolution of the oligosaccharide-binding pocket of plant IGPs. (A) Representative structures of IGP^ECD^ across different plant groups modeled by Alphafold 3. Ferns comprise both 12- and 13-LRR types. Orange and pink highlight the N-loopout motif and malectin domain, respectively. LRR regions are color-coded by plant groups. Cellooligomer-binding pocket residues are marked with sticks model. (B) Amino acid usage frequency of potential cellooligomer-binding site of each order. Order names are labelled in different font colors: angiosperms-green, gymnosperms-orange, ferns-purple, lycophytes-blue, and mosses-yellow. IGP numbers within each order are annotated in the bracket. The absence of the N-loopout motif is indicated short grey dash.

**fig. S15.**
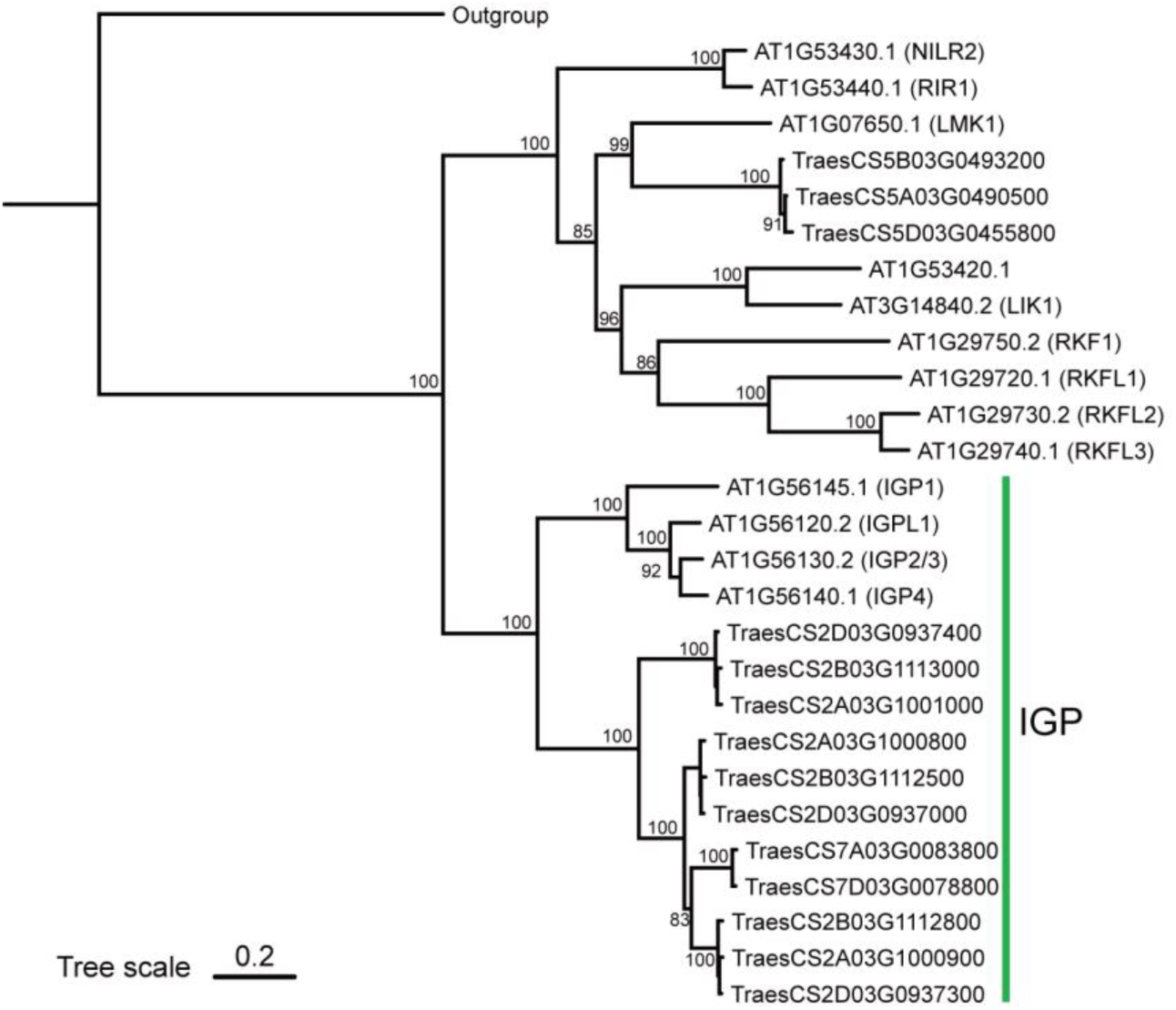
Phylogenetic analysis of wheat and Arabidopsis LRR-Malectin RLKs. Phylogenetic tree was constructed from 13 Arabidopsis LRR-Malectin RLKs and 14 wheat LRR-Malectin RLKs, using AT1G51880.3 as the outgroup.

**fig. S16.**
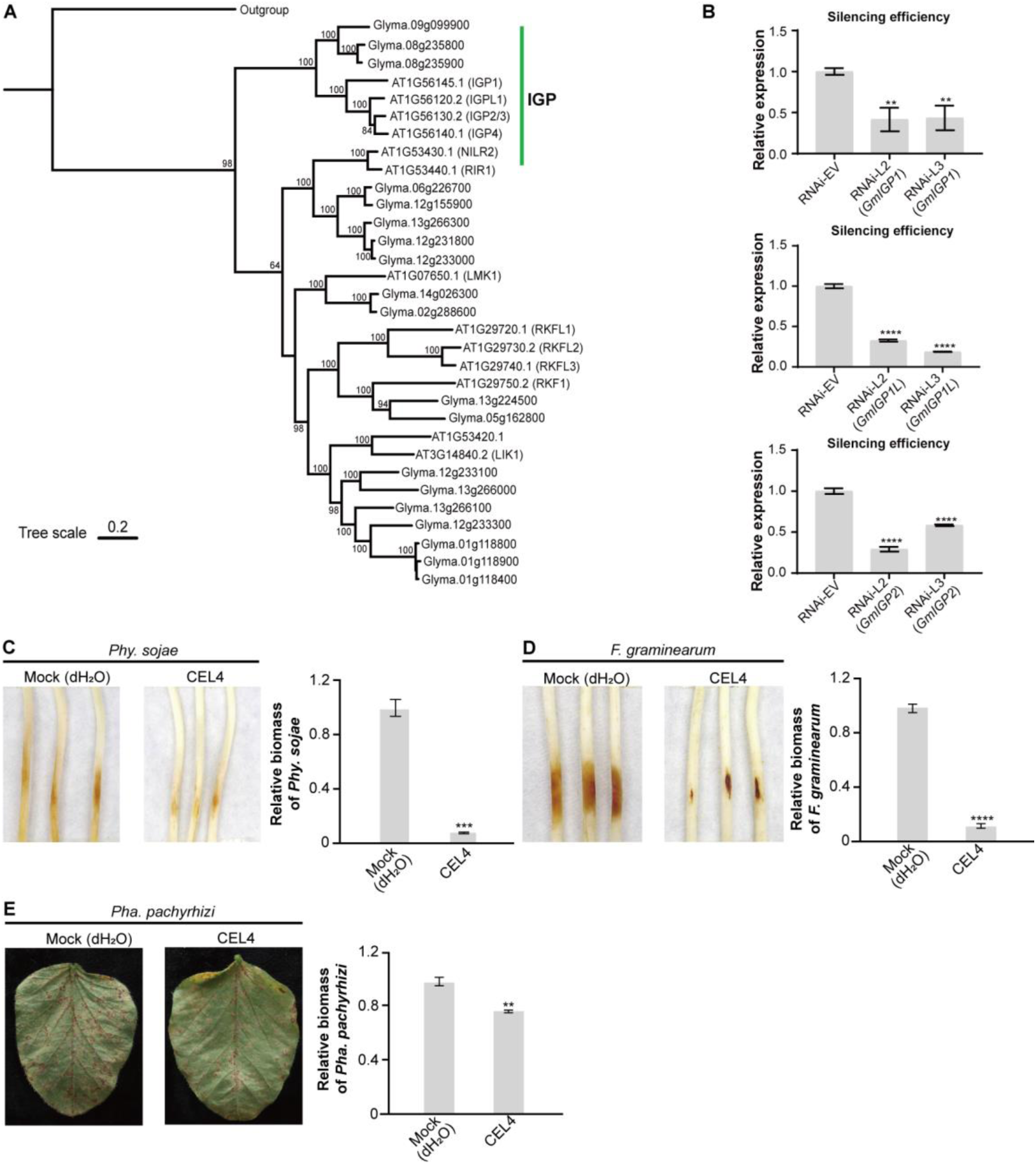
IGP-cellooligomer module confers broad-spectrum disease resistance in soybean. (**A**) Phylogenetic tree was constructed from 13 Arabidopsis LRR-Malectin RLKs and 19 *soybean* LRR-Malectin RLKs, using AT1G51880.3 as the outgroup. (**B**) The silencing efficiency of *GmIGP1*, *GmIGP1L*, and *GmIGP2* in soybean root hairs was quantified by qPCR in two independent *GmIGP*s co-silenced lines (RNAi-L2 and RNAi-L3), with *GmCYP2* serving as the reference gene for normalization. (**C**) Visual symptoms 48 hpi with *Phy. sojae* P6497 on soybean etiolated hypocotyls, which were pretreated separately with mock (dH_2_O) and CEL4 (10 μM). Representative photographs are shown (left). Relative biomass of *Phy. sojae* infecting etiolated soybean hypocotyls (at 48 hpi), as measured by genomic DNA quantitative PCR, and normalized to P6497 (right). (**D**) Visual symptoms 48 hpi with *F. graminearum* on soybean etiolated hypocotyls, which were pretreated separately with mock (dH_2_O) and CEL4 (10 μM). Representative photographs are shown (left). Relative biomass of *F. graminearum* infecting etiolated soybean hypocotyls (at 48 hpi), as measured by genomic DNA quantitative PCR, and normalized to *F. graminearum* (dH_2_O) (right). (**E**) Visual symptoms 7 dpi with *Pha. pachyrhizi* on soybean leaves, which were pretreated separately with mock (dH_2_O) and CEL4 (10 μM). Representative photographs are shown (left). Relative biomass of *Pha. pachyrhizi* infecting soybean leaves (at 7 dpi), as measured by genomic DNA quantitative PCR, and normalized to *Pha. pachyrhizi* (dH_2_O) (right).

**fig. S17.**
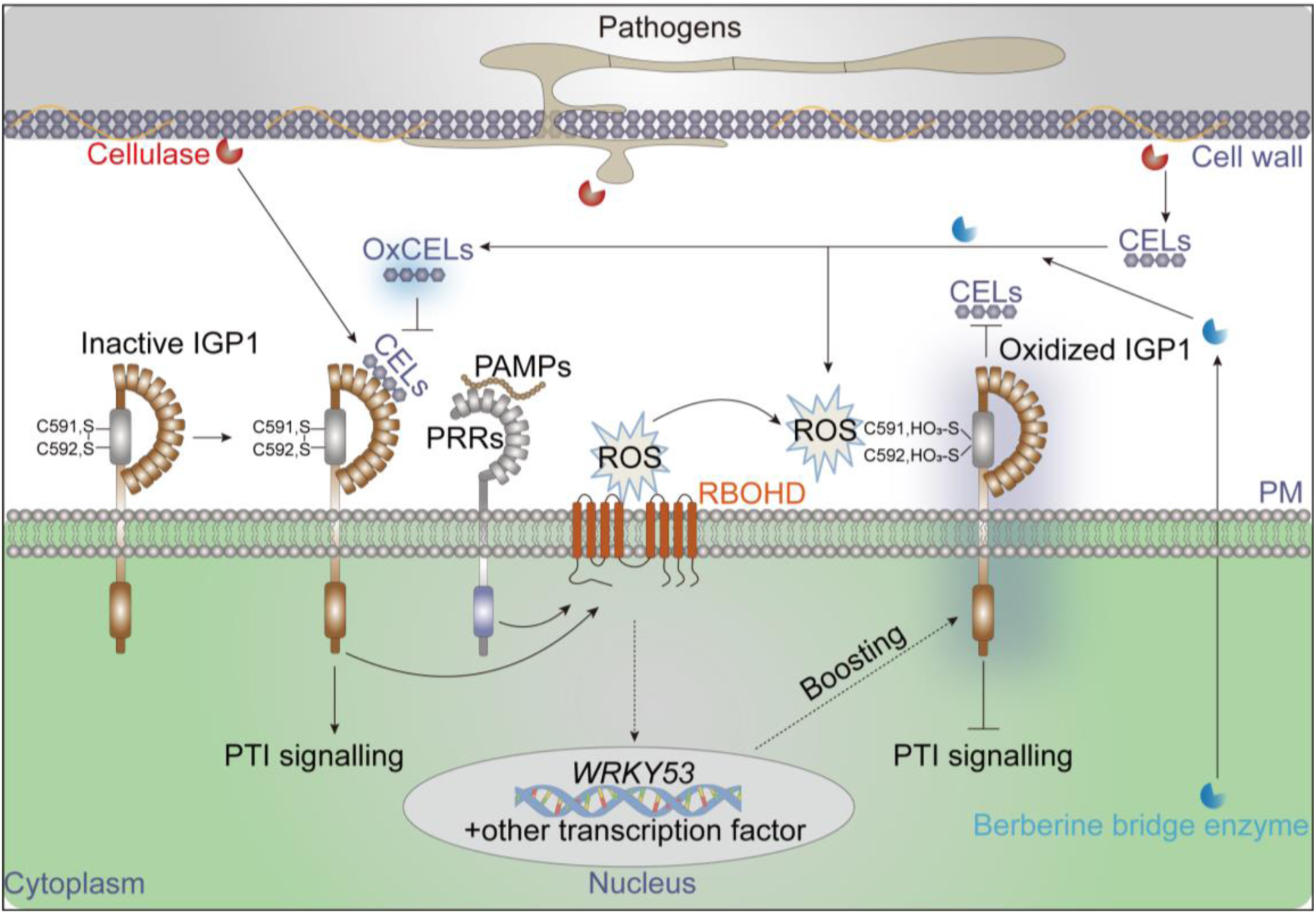
A model of IGP1-cellooligomer-mediated immune signaling. Plants fine-tune their immune homeostasis through the IGP1 receptor. Under resting conditions, IGP1 remains in an inactive state with low constitutive expression levels. Upon pathogen infection, secreted cellulases degrade cellulose in the plant cell wall, generating abundant cellooligomers (CELs) with DP ≥ 3. These oligosaccharides induce the upregulation of *IGPs* gene expression. Specifically, IGP1 binds to CELs via its extracellular 3rd to 5th leucine-rich repeat (LRR) domains, thereby activating basal immune responses such as reactive oxygen species (ROS) production, MAPK activation, and upregulation of defense-related genes. To maintain immune equilibrium, ROS-produced in response to CELs and other PAMPs-further enhance IGP1 protein accumulation through an unidentified transcriptional mechanism. Concurrently, these ROS oxidize the cysteine pair (C591-C592) within the extracellular malectin domain of IGP1. Oxidation of IGP1 reduces its binding capacity to CELs, thus preventing sustained immune activation. Additionally, plants employ Berberine bridge enzymes (BBEs) to oxidize CELs into OxCELs. This oxidation process serves a dual immunomodulatory function: OxCELs directly suppress CELs-induced immune responses, while the concomitant ROS production synergistically oxidizes IGP1, thereby collectively contributing to immune suppression. Collectively, these findings demonstrate that plants achieve precise regulation of immune homeostasis through redox-dependent modulation of both the IGP1 receptor and its ligands CELs.

**Table S1. Genomic data for 1,177 species.**

**Table S2. Transcriptome data for 97 species.**

**Table S3. Classification of LRR-Malectin RLKs based on phylogenetic inference.**

**Table S4. Mass spectrometry data for identified Cys oxidation on IGP1 ectodomain.**

**Table S5.**
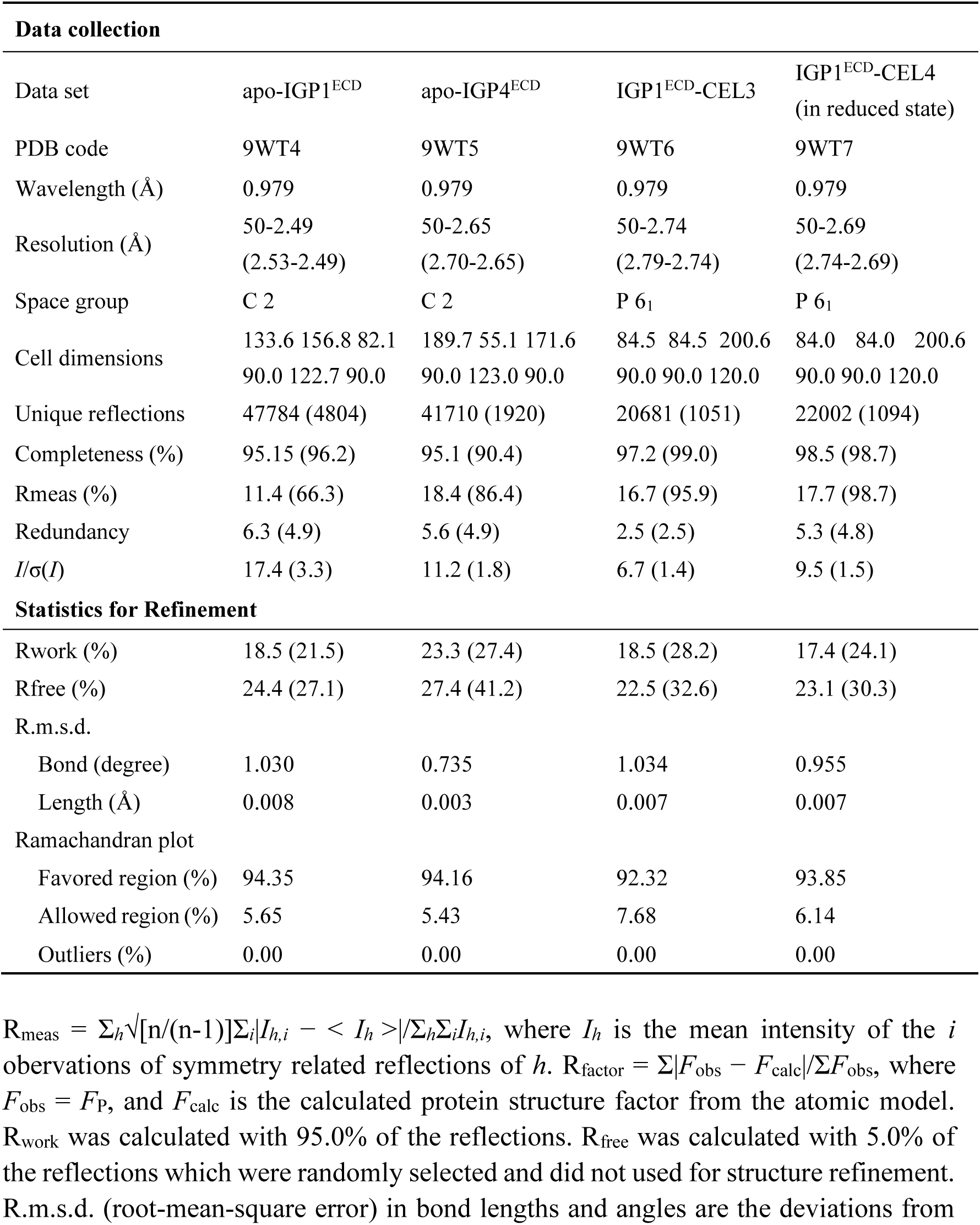
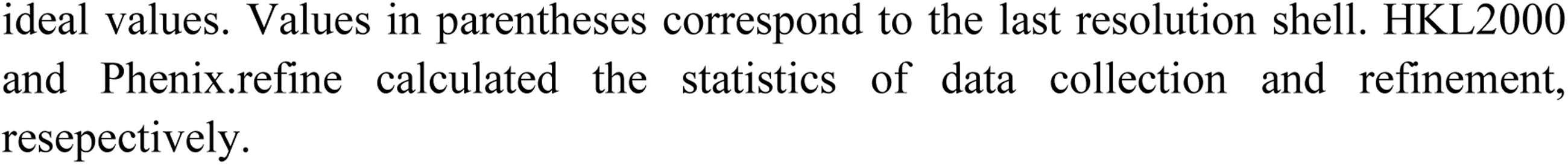
Summary of crystallography analyses.

| Data collection |  |  |  |  |
| --- | --- | --- | --- | --- |
| Data set | apo-IGP1<sup>ECD</sup> | apo-IGP4<sup>ECD</sup> | IGP1<sup>ECD</sup>-CEL3 | IGP1<sup>ECD</sup>-CEL4<br>(in reduced state) |
| PDB code | 9WT4 | 9WT5 | 9WT6 | 9WT7 |
| Wavelength (Å) | 0.979 | 0.979 | 0.979 | 0.979 |
| Resolution (Å) | 50-2.49<br>(2.53-2.49) | 50-2.65<br>(2.70-2.65) | 50-2.74<br>(2.79-2.74) | 50-2.69<br>(2.74-2.69) |
| Space group | C 2 | C 2 | P 6<sub>1</sub> | P 6<sub>1</sub> |
| Cell dimensions | 133.6 156.8 82.1<br>90.0 122.7 90.0 | 189.7 55.1 171.6<br>90.0 123.0 90.0 | 84.5 84.5 200.6<br>90.0 90.0 120.0 | 84.0 84.0 200.6<br>90.0 90.0 120.0 |
| Unique reflections | 47784 (4804) | 41710 (1920) | 20681 (1051) | 22002 (1094) |
| Completeness (%) | 95.15 (96.2) | 95.1 (90.4) | 97.2 (99.0) | 98.5 (98.7) |
| Rmeas (%) | 11.4 (66.3) | 18.4 (86.4) | 16.7 (95.9) | 17.7 (98.7) |
| Redundancy | 6.3 (4.9) | 5.6 (4.9) | 2.5 (2.5) | 5.3 (4.8) |
| *I*/$\sigma$(*I*) | 17.4 (3.3) | 11.2 (1.8) | 6.7 (1.4) | 9.5 (1.5) |
| Statistics for Refinement |  |  |  |  |
| Rwork (%) | 18.5 (21.5) | 23.3 (27.4) | 18.5 (28.2) | 17.4 (24.1) |
| Rfree (%) | 24.4 (27.1) | 27.4 (41.2) | 22.5 (32.6) | 23.1 (30.3) |
| R.m.s.d. |  |  |  |  |
| Bond (degree) | 1.030 | 0.735 | 1.034 | 0.955 |
| Length (Å) | 0.008 | 0.003 | 0.007 | 0.007 |
| Ramachandran plot |  |  |  |  |
| Favored region (%) | 94.35 | 94.16 | 92.32 | 93.85 |
| Allowed region (%) | 5.65 | 5.43 | 7.68 | 6.14 |
| Outliers (%) | 0.00 | 0.00 | 0.00 | 0.00 |

$R_{\text{meas}} = \sum_h \sqrt{[n/(n-1)] \sum_i |I_{h,i} - \langle I_h \rangle| / \sum_i I_{h,i}}$ , where $I_h$ is the mean intensity of the $i$ observations of symmetry related reflections of $h$ . $R_{\text{factor}} = \sum |F_{\text{obs}} - F_{\text{calc}}| / \sum F_{\text{obs}}$ , where $F_{\text{obs}} = F_P$ , and $F_{\text{calc}}$ is the calculated protein structure factor from the atomic model. $R_{\text{work}}$ was calculated with 95.0% of the reflections. $R_{\text{free}}$ was calculated with 5.0% of the reflections which were randomly selected and did not used for structure refinement. R.m.s.d. (root-mean-square error) in bond lengths and angles are the deviations from
ideal values. Values in parentheses correspond to the last resolution shell. HKL2000 and Phenix.refine calculated the statistics of data collection and refinement, respectively.

**Table S6. Structural features of IGPs across major plant groups.**

**Table S7. Natural mutation of CC motif from 321 structurally intact IGPs.**

